# An unconventional conductance is required for pacemaking of nigral dopamine neurons

**DOI:** 10.1101/2020.12.16.423073

**Authors:** Kevin Jehasse, Laurent Massotte, Sebastian Hartmann, Romain Vitello, Sofian Ringlet, Marie Vitello, Han Chow Chua, Stephan A. Pless, Dominique Engel, Jean-François Liégeois, Bernard Lakaye, Jochen Roeper, Vincent Seutin

**Affiliations:** Laboratory of Neurophysiology, GIGA-Neurosciences, University of Liège, Liège, Belgium; Institute for Neurophysiology, Goethe University, Frankfurt, Germany; Laboratory of Medicinal Chemistry, CIRM, University of Liège, Liège, Belgium; Department of Drug Design and Pharmacology, University of Copenhagen, Denmark; Laboratory of Molecular Regulation of Neurogenesis, GIGA-Stem Cells, University of Liège, Liège, Belgium

**Author notes:** Both last authors contributed equally to this work (Jochen Roeper, Vincent Seutin). Correspondence: Vincent Seutin (Laboratory of Neurophysiology, GIGA-Neurosciences, University of Liège).

## Abstract

Although several ionic mechanisms are known to control rate and regularity of the pacemaker in dopamine (DA) neurons from the substantia nigra pars compacta (SNc), a conductance essential for pacing has yet to be defined. Here we provide pharmacological evidence that pacemaking of SNc DA neurons is enabled by an unconventional conductance. We found that 1-(2,4-xylyl)guanidine (XG), an established blocker of gating pore currents in mutant voltage gated sodium channels, selectively stops pacemaking of DA SNc neurons and is without effect on the main pore of their voltage-gated channels. We isolated a voltage-dependent, non-inactivating XG-sensitive current of 20-25 pA which operates in the relevant subthreshold range and is carried by both Na^+^ and Cl^-^ ions. While the molecular identity of this conductance remains to be determined, we show that this XG-sensitive current is crucial to sustain pacemaking in these neurons.

## INTRODUCTION

Midbrain dopamine (DA) neurons are slow pacemaker neurons. They are involved in diverse psychomotor functions including control of movement, reward, motivation and cognition (Nieoullon, 2002; Schultz, 2007; Wise, 2004). Although several subpopulations of midbrain DA neurons have been described based on their projections, transcriptomic profiles and electrophysiological properties (Evans et al., 2017; Farassat et al., 2019; Liss and Roeper, 2008; Margolis et al., 2008; Neuhoff et al., 2002; Philippart et al., 2016; Poulin et al., 2014), the majority of DA neurons are able to fire action potentials (AP) spontaneously at low frequency, in the range of 1 to 5 Hz, which might ensure a basal level of DA signaling in the target areas (Albin et al., 1989; Gonon and Bloch, 1998; Grace and Bunney, 1984). They are also known to switch their firing mode from pacemaking to burst activity in response to synaptic input (Grace and Bunney, 1984).

While the onset of bursting is clearly linked to changes in synaptic inputs involving activation of NMDA receptors (Blythe et al., 2007; Destreel et al., 2019; Galtieri et al., 2017; Johnson et al., 1992), the exact mechanism(s) that enable DA neuron slow pacemaking is still unknown. Blocking the hyperpolarization-activated cation channels (HCN) sustaining *I_h_* does not stop the autonomous activity of DA neurons, but it slows the firing frequency of some of them (Neuhoff et al., 2002; Seutin et al., 2001). It was also suggested that Ca^2+^ drives the spontaneous firing through L-type voltage-gated calcium (Ca_v_) channels, as applying dihydropyridines (DHPs) slows or even silences the pacemaking of DA SNc neurons (Chan et al., 2007; Mercuri et al., 1994; Nedergaard et al., 1993; Puopolo et al., 2007). However, these experiments were made using high concentration of DHPs, leading to possible off-target actions or these effects were observed only in a subset of the recorded cells (Puopolo et al., 2007). Another study found no effect of lower concentrations of DHPs (Guzman et al., 2009).

Understanding what happens during the interspike interval (ISI) is key to uncover the nature of the autonomous activity of DA neurons. The coefficient of variation of the interspike interval (ISI) in DA SNc neurons *ex vivo* is very low, in the range of 5% (de Vrind et al., 2016). The current amplitude during the ISI has been estimated to be around 5 pA (Khaliq and Bean, 2008). Taking into account the gating properties and the single channel conductance of most voltage-gated ion channels, which lies around 20 pS, this would mean that few ion channels would underlie this current. Given the stochastic nature of ion channel opening, such a current would inevitably be very noisy, which is incompatible with the small coefficient of variation of the ISI of DA neurons. A second option would be the result from the almost perfect cancellation of inward and outward currents ensuring the robustness of pacemaking (O’Leary et al., 2014). In DA neurons, it could be mediated by fast activating K^+^ channels K_v_4 (sustaining *I_A_*) cooperating with HCN channels, but blocking both channels only affects their rebound firing (Amendola et al., 2012).

A third possibility would be the recruitment of many small conductance pores to ensure a steady pacemaking. We therefore turned our attention to unconventional pores displaying a very low conductance, in the range of fS, such as currents generated from gating pores (“gating pore currents” or “omega currents” (*I_ω_*)). *I_ω_* is generally the result of a pathological mutation in the voltage sensor domain, creating a pore through which ions can flow (Jiang et al., 2018; Sokolov et al., 2007; Tombola et al., 2005). Their conductance has been estimated to be 1-6% the conductance of the main α pore of voltage-gated channels (Held et al., 2016). It was subsequently found that *I_ω_* generated from a mutant voltage-gated sodium (Na_v_) channel is selectively blocked by 1-(2,4-xylyl)guanidine (XG) (Sokolov et al., 2010). Since XG does not interact with the α pore of Nav channels, we used it as a pharmacological tool to probe for such currents in DA SNc neurons.

## RESULTS

### XG inhibits the slow pacemaker activity of DA neurons across the entire midbrain

We first performed extracellular recordings on acute brain slices from adult rats to investigate the effect of XG on the spontaneous activity on DA neurons. We applied synaptic blockers (10 μM CNQX, 1 μM MK-801, 1 μM CGP55845, 10 μM SR9551 and 1 μM sulpiride) before and during the superfusion of XG to prevent any presynaptic effect of the compound. We observed that application of 2 mM XG stops the firing of these neurons (fig. 1a). We then tested lower concentrations of XG (fig. 1b) and found that the compound inhibits the spontaneous firing of DA neurons in a concentration-dependent manner with an IC_50_ of 486 ±51 μM (n = 6 for each concentration; fig. 1c) and a Hill slope of 1.287 ± 0.194. This result is consistent with the pharmacology of *I_ω_* (Sokolov et al., 2010) and indicates that the inhibition is probably due to the interaction of XG with a single target. Since the carbonate salt of XG, which was used in these experiments, is difficult to solubilize at physiological pH, we synthetized a mesylate salt of XG which is much easier to use (see methods). We performed the same experiments with this compound and obtained an IC_50_ of 427 ± 43 μM, which is not significantly different from the one of the carbonate salt (n = 6 for each condition, p = 0.71, extra sum-of-squares F test; fig. 1c). The Hill slope was 1.273 ± 0.153. Interestingly, we obtained similar results on DA neurons located in the ventral tegmental area (VTA) using the mesylate salt of XG. These experiments yielded an IC_50_ for the blocker of 332.5 ± 95 μM and a Hill slope of 1.052 ± 0.282 (n = 6 for each condition, p = 0.15, extra sum-of-squares F test; fig. 1c).

**Figure 1.**
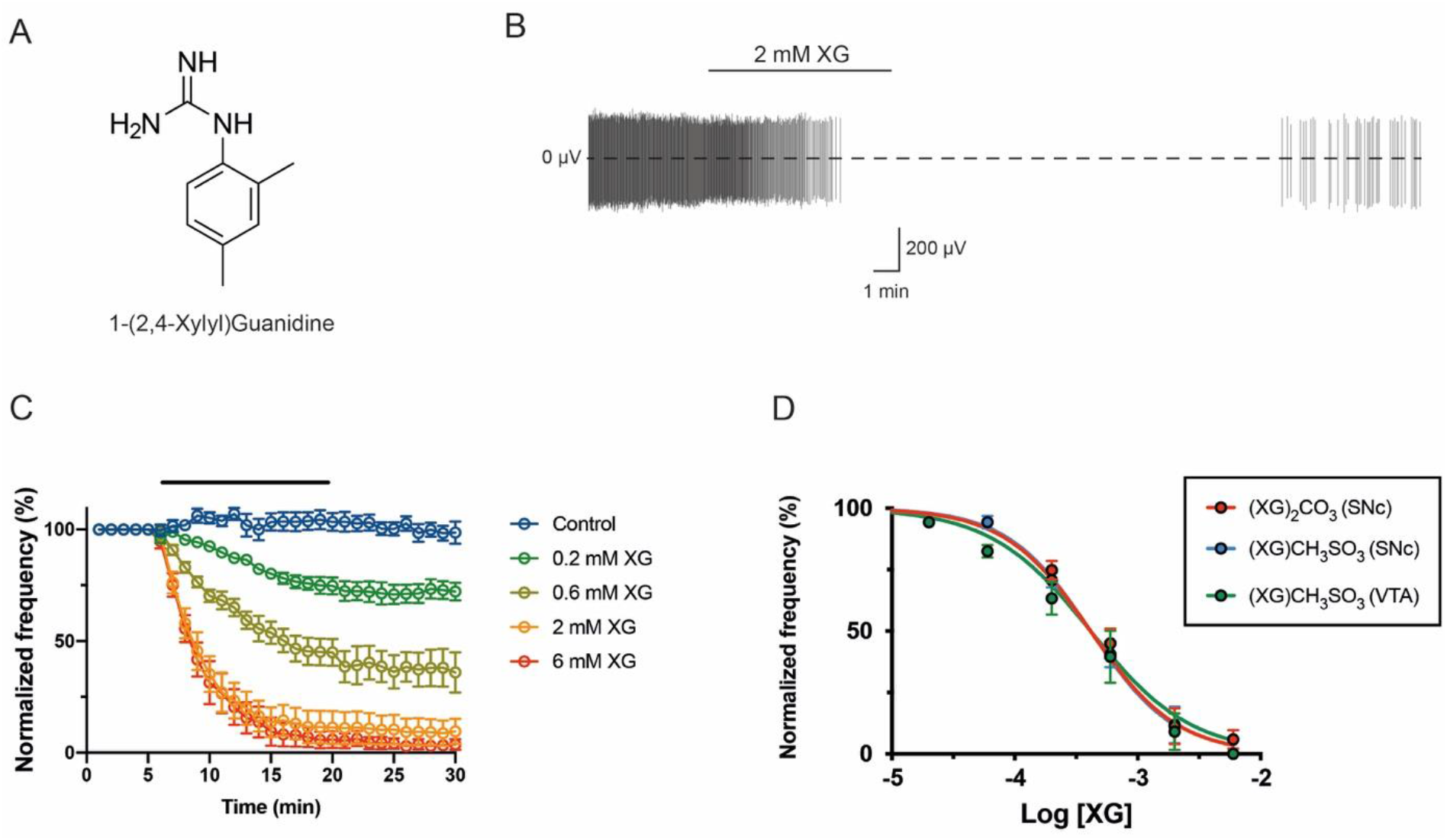
XG inhibits the spontaneous activity of rat DA SNc neurons. **A.** Molecular structure of XG. **B.** Representative extracellular recording of the spontaneous firing of a single DA SNc neuron. The continuous line corresponds to the time of application of XG on the slice. Note that the effect of XG is partially reversible after a long washout. **C.** Summary data of the evolution of the firing frequency according to the time of exposition and the concentration. Each point corresponds to the mean normalized frequency ± SEM (in %, n=6). The black line represents the time during which the tested solution was superfused. **D.** Concentration-response curve of the inhibitory effect of XG. All values were normalized to the control firing frequency. Note that the potency is very similar for the two different salts and in the two regions tested (n=6 for each condition).

We also carried out cell-attached recordings on synaptically isolated mouse DA SNc neurons to verify whether the pacemaker inhibition by XG is species-dependent. We applied 2 mM XG after a stable baseline of spontaneous discharge was established (fig. 2a, b). After > 10 min, we opened the cells to the whole-cell configuration in order to fill the neuron with neurobiotin for post-hoc immunofluorescence identification (fig. 2e). We observed a strong reduction of firing frequency of 1.48 ± 0.16 Hz (n = 10, p < 0.01, t-test; fig. 2c) across the entire SNc (fig. 2d). These results indicate that the pacemaker inhibition by XG in DA SNc neurons is similar in both rats and mice.

**Figure 2.**
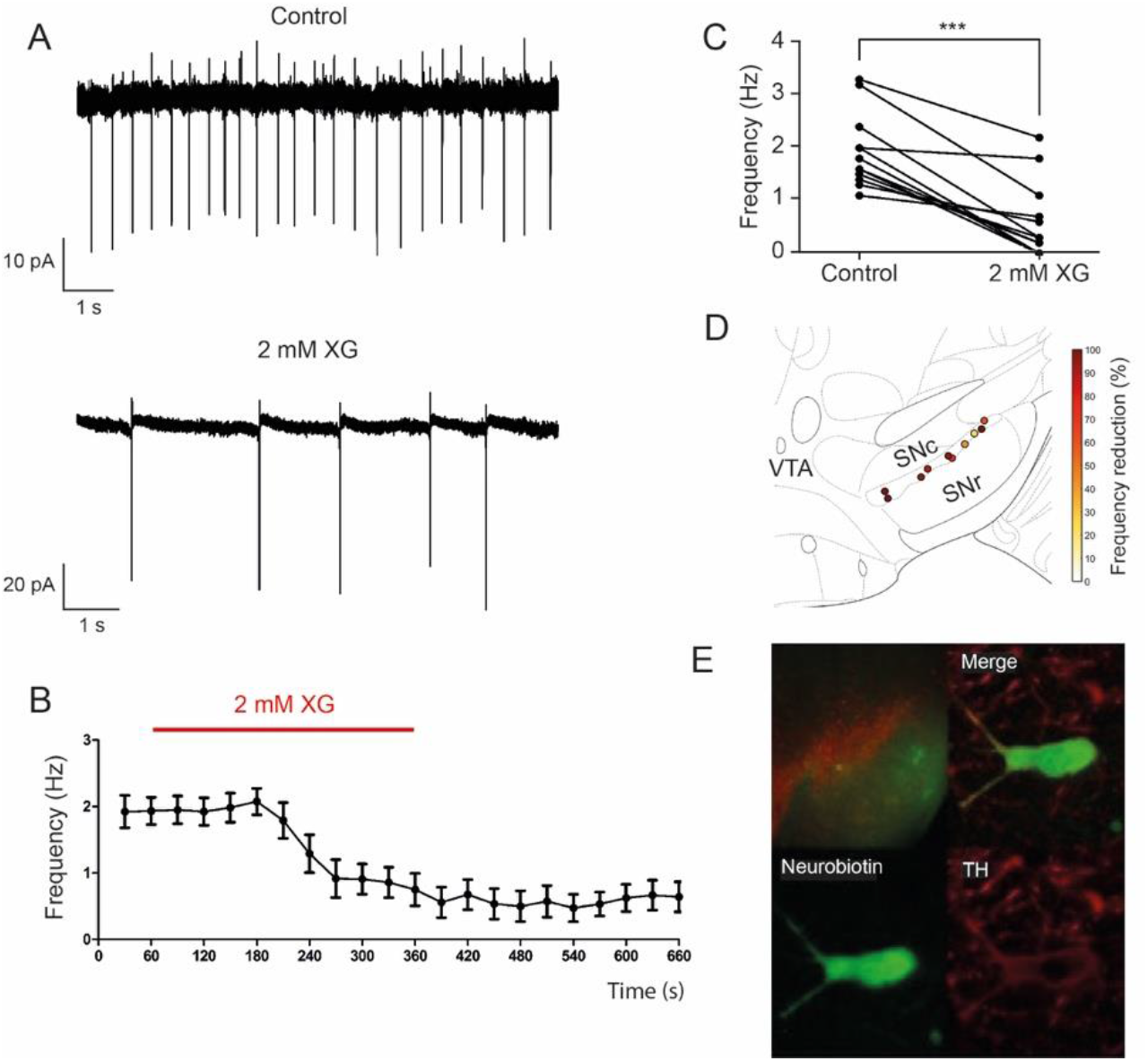
XG inhibits the spontaneous activity of mouse DA SNc neurons. **A.** Representative cell-attached recording showing the inhibitory effect of 2 mM XG. **B.** Group data showing the average inhibition produced by XG over time (n=10). **C.** Summary data of individual experiments showing a reduction of the firing frequency (n=10). **D.** Heat map of the inhibitory effect of XG according to the localization of the recorded neurons within SNc. **E.** *Post-hoc* identification of a recorded neurons filled with neurobiotin as being DA based on TH immunofluorescence.

### XG selectively inhibits the pacemaker activity of DA SNc neurons

We next investigated the mechanism of inhibition with whole-cell patch-clamp recordings. SNc DA neurons were identified as described in the methods (Fig. S1). In line with our previous experiments, extracellular application of XG was able to stop the spontaneous activity of DA SNc neurons (1.94 ± 0.18 Hz in control vs 0.02 ± 0.001 Hz in 2 mM XG, n = 8, p = 0.0078, Wilcoxon test; fig. 3a-b). Intriguingly, intracellular application of XG also inhibited the pacemaking (1.91 ± 0.29 Hz in control vs 0.0003 ± 0.0003 Hz during intracellular application of 2 mM XG, n = 6, p = 0.0312, Wilcoxon test; fig. 3c). We also tested 2 mM XG on GABA substantia nigra pars reticulata neurons, which are fast pacemaker neurons. XG did not significantly change the firing frequency of these neurons (8.18 ± 1.07 Hz in control vs 8.52 ± 1.06 Hz in 2 mM XG, n = 7; p = 0.21, t-test; fig. 3f-g).

**Figure 3.**
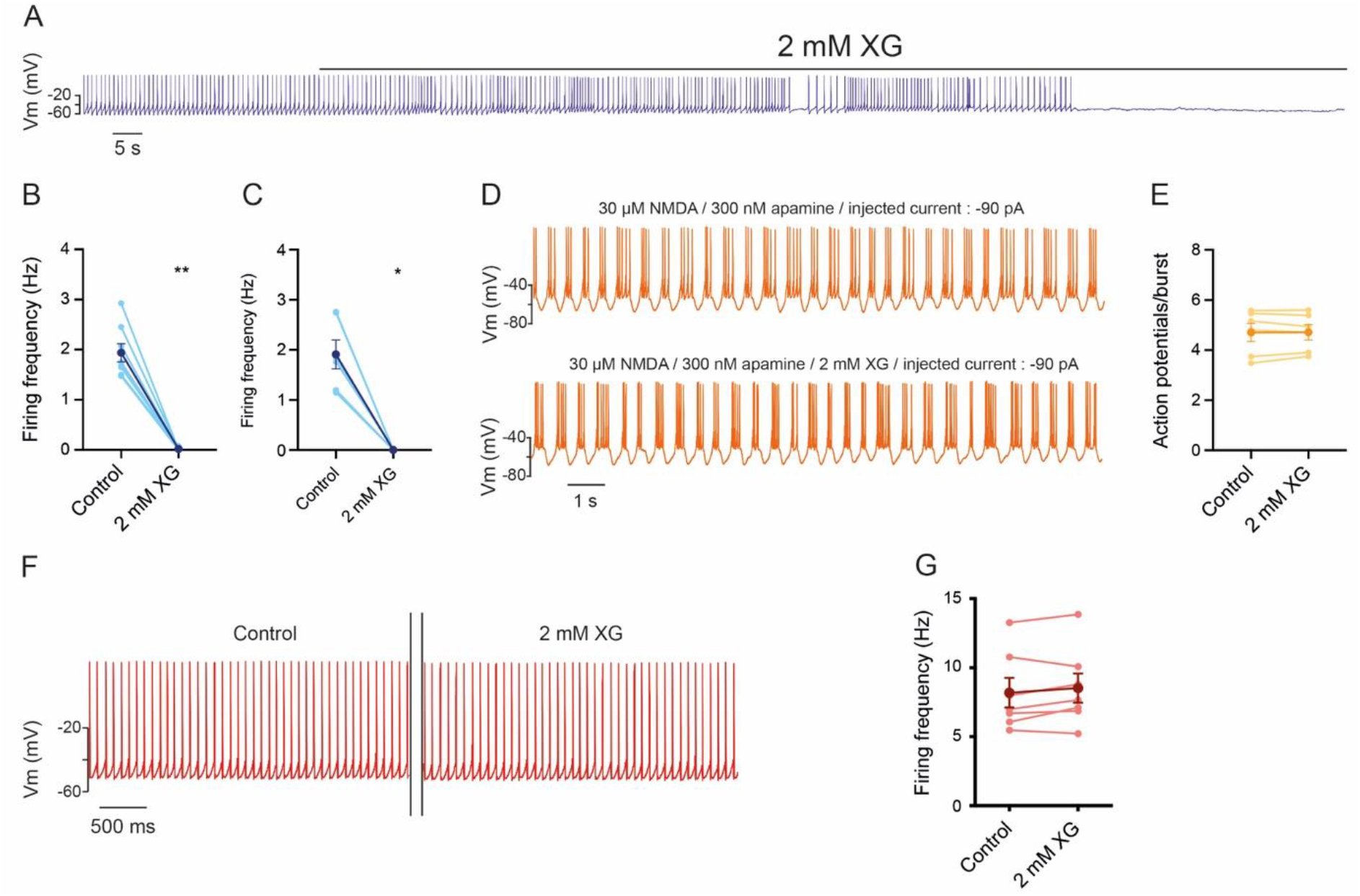
XG selectively inhibits the pacemaking activity of DA SNc neurons. **A**. Representative recording of a single DA SNc neuron. The black line corresponds to the application of 2 mM XG on the slice. **B.** Summary data of the effect of the extracellular application of 2 mM XG on the firing frequency. The light blue data correspond to individual neurons (n = 8) and the dark blue correspond to the mean ± SEM of all recorded neurons. XG abolished the firing (Wilcoxon test, **p < 0.01). **C.** Summary data of the effect of intracellular application of 2 mM XG on the firing frequency. The light blue data corresponds to individual neurons (n = 6) and the dark blue corresponds to the mean ± SEM of all recorded neurons. XG abolished the firing (Wilcoxon, *p < 0.05). **D.** Representative recording of bursting activity in a single DA SNc neuron. **E.** Summary data of the effect of application of 2 mM XG on the number of APs per burst. The light orange data corresponds to individual neurons (n = 6) and the darker orange corresponds to the mean with SEM of all recorded neurons. XG did not significantly affect the number of APs/burst (t-test, p = 0.9). **F.** Representative recording of a fast firing GABAergic SNr neuron. **G.** Summary data of the effect of the extracellular application of 2 mM XG on the firing frequency of SNr neurons. The light red data corresponds to individual neurons (n = 7) and the dark red corresponds to the mean ± SEM of all recorded neurons. XG did not affect the firing frequency in these neurons (t-test, p = 0.21).

Importantly, the membrane potential of DA SNc neurons during XG application stayed between −50 and −55 mV, indicating that the inhibition is not due to a strong hyperpolarization of the neurons. This was confirmed in voltage clamp experiments, in which we clamped the cells at −50 mV and imposed a hyperpolarizing step to −60 mV. The whole-cell conductance was not affected by 2 mM XG (1.98 ± 0.23 nS in control, 1.97 ± 0.23 nS in the presence of the drug and 2.06 ± 0.23 nS during washout, n = 8 for each condition, p = 0.17, ANOVA-1; fig. S2). The holding current at – 60 mV was unaffected as well. Taken together, these experiments demonstrate that XG inhibits the slow pacemaking of DA neurons specifically, reproducibly and without opening ion channels. In addition, since two different salts yielded similar results, we conclude that the xylylguanidinium moiety is responsible for the observed effect.

Injection of a small positive current in the range of 10-30 pA was able to reinstate action potential firing, indicating that the inhibition is not due to blockade of the main pore of Na_v_ channels (fig. 4a). We performed current injections from 0 pA to 50 pA for 2 s to analyze three parameters of the AP: threshold, half-width and maximal slope. Application of 2 mM XG did not change the threshold (p = 0.88), the half-width (p = 0.30) or the maximal slope (p = 0.40) of the AP whatever the current amplitude (n = 5 for all conditions, ANOVA-2; fig. 4b, Table 1). We also performed 20 pA steps from 0 pA to 160 pA for 2 s to determine the shift in current needed to generate pacemaking. The linear regression (fig. 4c) of individual frequency-current curves shows that there was a shift from −19.43 ± 7.56 pA in control to 4.46 ± 2.32 pA in 2 mM XG (n = 6, p = 0.03, t-test; fig. 4e), indicating that the amplitude of the blocked current is −23.89 ± 8.21 pA. We also did not observe a significant change in the gain (68.63 ± 13.69 Hz/nA in control vs 84.79 ± 11.60 Hz/nA in 2 mM XG, n = 6, p = 0.23, t-test; fig. 4d), which is consistent with the results from the characterization of the action potential parameters. These results demonstrate that the inhibition by XG is not due to the modulation of voltage-gated channels involved in AP generation but rather involves the blockade of a small but essential inward current in the subthreshold range.

**Figure 4.**
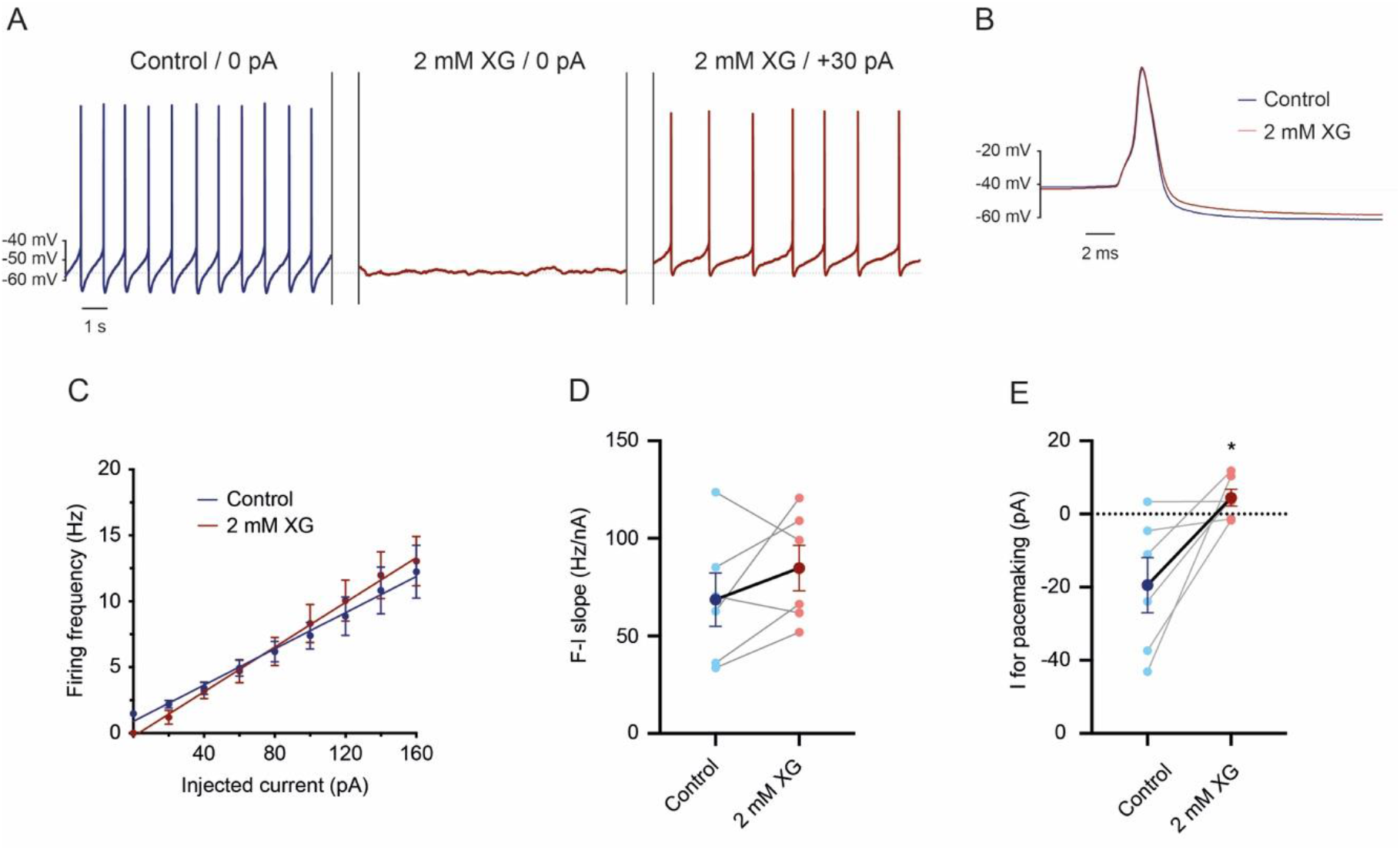
XG does not modulate voltage-gated channels involved in the action potential. **A.** Representative recording showing that the firing of a DA neuron can be recovered in XG upon injection of a small positive current. **B.** Superposition of one action potential in control conditions and in XG with a +30 pA current. **C.** Mean frequency-current relationship of 6 DA neurons in control conditions (blue) and in the presence of 2 mM XG (red). The frequency was measured using 20 pA steps from 0 pA to 160 pA for 2 s. Only the three first APs were used to calculate the firing frequency. **D.** Summary data of the effect of 2 mM XG on the gain (F-I slope). The individual slopes of the linear regression from single cells are represented in light color (n = 6 for both conditions) while the means ± SEM are represented in darker color. **E.** Summary data of the effect of 2 mM XG on the minimal current needed to generate spontaneous activity. The individual minimal current for pacemaking to occur are represented in light color (n = 6 for both conditions) while the means ± SEM are represented in darker color. The current was determined based on the current value at which the linear regression intercepts with the current axis. The amplitude of the current blocked by 2 mM XG is 23.9 ± 8.2 pA.

**Table 1.** Summary results of the half-width, maximal slope and threshold of action potentials before and after application of XG. XG does not affect any of the three parameters.

|  | 0 pA |  | 10 pA |  | 20 pA |  | 30 pA |  | 40 pA |  | 50 pA |  |
| --- | --- | --- | --- | --- | --- | --- | --- | --- | --- | --- | --- | --- |
|  | Control | XG | Control | XG | Control | XG | Control | XG | Control | XG | Control | XG |
| Half-width (ms) | 2.00 ± 0.25 | - | 1.98 ± 0.15 | 1.98 ± 0.19 | 1.96 ± 0.15 | 2.17 ± 0.17 | 2.13 ± 0.2 | 2.23 ± 0.17 | 2.24 ± 0.16 | 2.33 ± 0.13 | 2.14 ± 0.21 | 2.32 ± 0.16 |
|  | n = 5 |  | n = 5 | n = 4 | n = 5 | n = 5 | n = 5 | n = 5 | n = 5 | n = 5 | n = 5 | n = 5 |
| Max slope (ms) | 83.44 ± 10.47 | - | 79.04 ± 8.33 | 92.12 ± 11.38 | 74.40 ± 7.49 | 78.67 ± 11.87 | 69.79 ± 7.04 | 74.34 ± 10.88 | 60.00 ± 7.38 | 68.97 ± 9.82 | 65.31 ± 5.16 | 58.17 ± 7.33 |
|  | n = 5 |  | n = 5 | n = 4 | n = 5 | n = 5 | n = 5 | n = 5 | n = 5 | n = 5 | n = 5 | n = 5 |
| Threshold (mV) | -35.13 ± 1.90 | - | -33.86 ± 2.34 | -35.64 ± 2.71 | -34.05 ± 1.98 | -33.32 ± 2.54 | -32.65 ± 2.13 | -32.46 ± 2.91 | -31.57 ± 2.43 | -32.53 ± 1.83 | -32.62 ± 1.81 | -39.68 ± 2.01 |
|  | n = 5 |  | n = 5 | n = 4 | n = 5 | n = 5 | n = 5 | n = 5 | n = 5 | n = 5 | n = 5 | n = 5 |

Since DA neurons display two firing modes *in vivo,* we induced bursting activity by superfusing 30 μM NMDA and 300 nM apamin to block SK channels, while injecting constant hyperpolarizing current (Destreel et al., 2019; Johnson et al., 1992) to test whether XG could modulate this firing mode. 2 mM XG did not modify the bursting: thus the number of bursts per second and the number of APs per burst did not change during the whole recording time (4.71 ± 0.36 APs in control vs 4.72 ± 0.31 in 2 mM XG, n = 6, p = 0.9, t-test; fig. 3d-e). These observations demonstrate that XG selectively blocks pacemaking without interfering with other types of firing patterns such as bursting activity (at least in this *ex vivo* model) in these neurons. This further demonstrates the very good selectivity of the drug on pacemaking of DA neurons.

### XG-sensitive conductance is crucial for oscillatory events and pacemaking in DA SNc neurons

In DA neurons, the spontaneous firing involves a combination of different conductances, as blocking only Nav or Cav is not sufficient to fully block autonomous activity (Drion et al., 2011; Guzman et al., 2009; Kang and Kitai, 1993). On the contrary, in all our whole-cell recordings, application of 2 mM XG was enough to silence DA neuron activity (fig. 2a). We next investigated whether these XG-sensitive channels can facilitate different oscillatory behaviors in DA neurons. We first studied autonomous oscillations by applying 1 μM TTX (fig. 5b). As previously shown by others (Guzman et al., 2009; Sun et al., 2017), we found that these oscillations are blocked by 5 μM nifedipine, suggesting that they are mediated by L-type Cav channels (fig. 5a). Application of 2 mM XG abolished these oscillations (n = 6; fig. 5b). We then investigated whether XG could still inhibit pacemaker activity in the absence of L-type Cav-mediated oscillations. We applied 5 μM nifedipine to block the latter. This did not change the firing frequency (1.96 ± 0.22 Hz in control vs 1.79 ± 0.18 Hz in 5 μM nifedipine, n = 6, p = 0.25, Friedman test; fig. 5c,d). These results are consistent with the literature (Drion et al., 2011; Guzman et al., 2009). Application of 2 mM XG also led to an almost complete inhibition of the spontaneous activity of DA SNc neurons (0.05 ± 0.03 Hz in 5 μM nifedipine + 2 mM XG vs 1.78 ± 0.18 Hz in 5 μM nifedipine only, p = 0.0433, Friedman test; fig. 5c, d). These data strongly suggest that an XG-sensitive current *(I_XG_)* is crucial to recruit various types of depolarizing channels to drive oscillatory events in DA SNc neurons.

**Figure 5.**
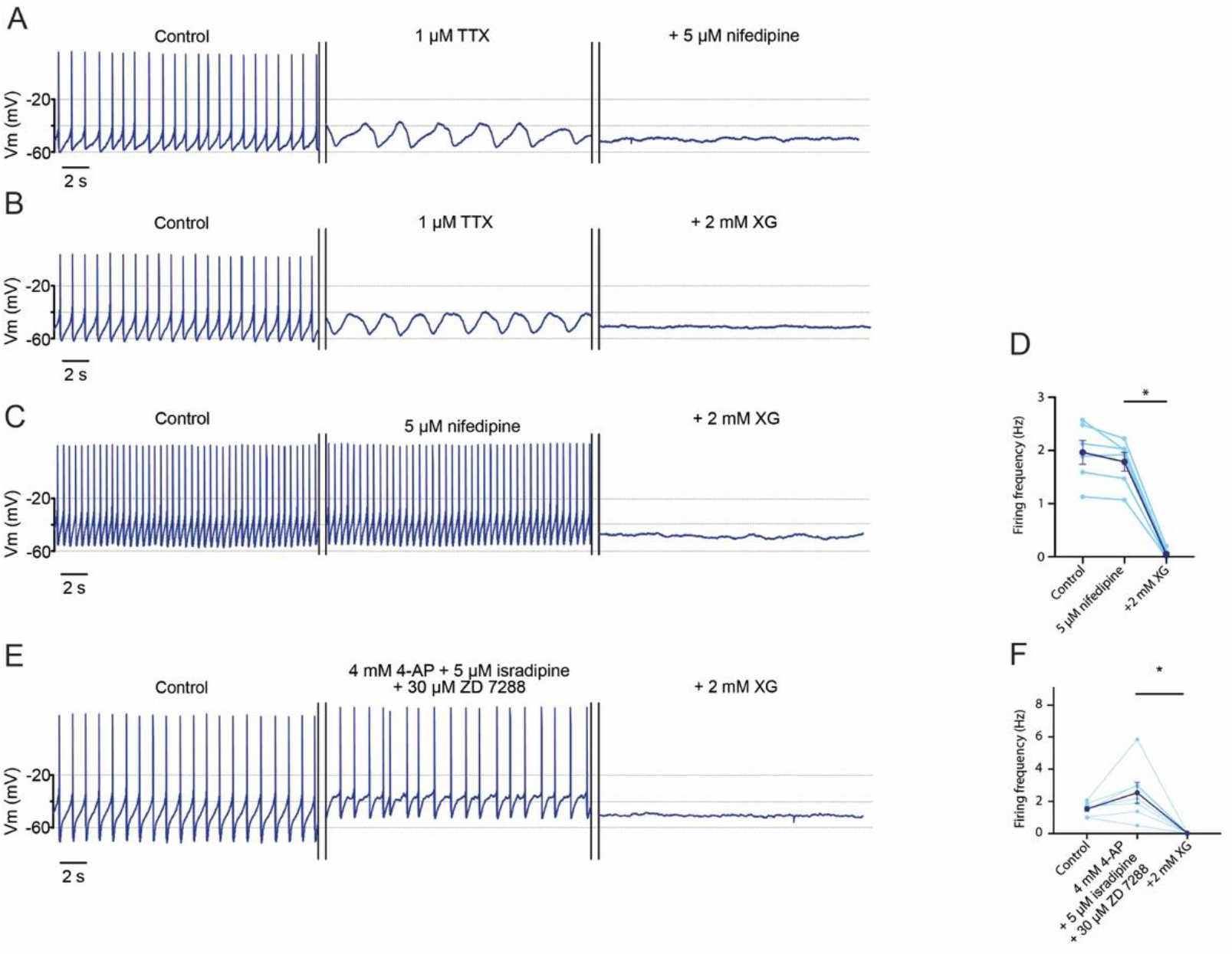
XG-sensitive conductance are essential to various oscillatory phenomena in DA SNc neurons. **A.** Applying 1 μM TTX on a spontaneous active DA neuron reveals oscillations of membrane potential (middle panel) that are nifedipine-sensitive (right panel), suggesting that they are mediated by L-type Cav channels. **B.** Representative recording of the effect of 2 mM XG on membrane potential oscillations. Application of 2 mM XG was able to silence them. **C.** Representative recording of the effect of 2 mM XG while Ca^2+^ oscillations are blocked by 5 μM nifedipine. In these conditions, the spontaneous firing is maintained (middle panel) and is still sensitive to 2 mM XG (right panel). **D.** Summary data of the effect of 2 mM XG on the firing frequency while Ca^2+^ oscillations are blocked. The light blue data corresponds to individual neurons (n = 6) and the dark blue corresponds to the mean ± SEM of all recorded neurons. **E.** Representative recording of the effect of 2 mM XG on the pacemaking activity while Kv4, Ca_v_1.3 and HCN channels are blocked. When the spontaneous activity remains in the presence of 4 mM 4-AP, 5 μM isradipine and 30 μM ZD 7288 (middle panel), application of 2 mM XG is still able to silence the neurons (right panel). **F.** Summary data of the effect of 2 mM XG on the firing frequency while K_v_4, Ca_v_1.3 and HCN channels are blocked. The light blue data corresponds to individual neurons (n = 6) and the dark blue corresponds to the mean ± SEM of all recorded neurons.

Given that the robustness of the pacemaking activity in DA neurons may involve a cooperation between Na_v_, K_v_4, Ca_v_1.3 and HCN channels, we next investigated whether blocking the all three categories of channels would impact on the spontaneous activity of DA SNc and whether XG would still silence them. Application of 4 mM 4-aminopyridine (4-AP), 5 μM isradipine and 30 μM ZD 7288 to respectively block Kv4, Cav1.3 and HCN channels did not change the firing frequency (1.53 ± 0.15 Hz in control vs 2.54 ± 0.64 Hz in 4 mM 4-AP, 5 μM isradipine and 30 μM ZD 7288, n = 6 for each condition, p = 0.10, ANOVA-1, fig. 5e,f) while adding 2 mM XG to the blockers significantly abolished the firing (0.02 ± 0.01 Hz in blockers + 2 mM XG, vs blockers, n = 6 for each condition, p = 0.02, ANOVA-1, fig. 5e,f). Taken together, these results strengthen the hypothesis that *I_XG_* is driving the spontaneous activity of DA neurons rather than promoting a cooperation between different conductances.

NALCN channels have been suggested to play a role in the pacemaking of DA neurons (Khaliq and Bean, 2010; Philippart and Khaliq, 2018). To test this hypothesis, we first performed extracellular recordings using 100 μM Gd^3+^, a known inhibitor of NALCN channels (Chua et al., 2020). This agent did not significantly alter the firing frequency of DA neurons (fig. S3a). As block by Gd^3+^ is not specific to NALCN channels (Boone et al., 2014), we tested the possibility of a direct effect of XG on NALCN. To this end, we heterologously expressed the NALCN complex (NALCN, UNC80, UNC79 and FAM155A) in *Xenopus laevis* oocytes and performed two-electrode voltage-clamp experiments. We only observed minimal inhibition of NALCN currents by XG (37 ± 3.9 % inhibition at −100 mV for 3 mM XG, n = 6), while 10 μM Gd^3+^ potently inhibited NALCN currents (74 ± 9.3 % inhibition at −100 mV, n = 6) (fig. S3b). Therefore, the effect of XG on pacemaking is unlikely to be due to inhibition of NALCN channels.

### No clear evidence for *I_ω_* in DA SNc neurons

Our data so far suggests that XG inhibits an as yet unidentified channel essential for pacemaking in DA SNc neurons. As XG blocks voltage-sensor-related ω pores but not the conventional α pores in mutant Na_v_ channels (Sokolov et al., 2010), one possibility is that DA SNc neurons express a yet to be defined channel protein that provides a physiological XG-sensitive ω pore for pacemaking. ω pores are often created by the replacement of an arginine in the fourth segments (S4) of the voltage sensor domain of mutant voltage-gated channels (Sokolov et al., 2007; Tombola et al., 2005). Depending on the location of the missing arginine, it generates either a small inward or outward current at hyperpolarized or depolarized potentials respectively (Held et al., 2016; Jiang et al., 2018). Therefore, we next aimed to obtain evidence for an involvement of *I_ω_* and to identify a molecular candidate potentially coding for such a current in DA SNc neurons.

First, we used voltage-clamp recordings to isolate *I_XG_* and examine its shape. Indeed, it had previously been shown that inward *I_ω_* is prominent at negative (−150 to −100 mV) potentials and becomes smaller at less negative (−50 mV) voltages (Moreau et al., 2014; Sokolov et al., 2010). We performed ramp recordings, from −100 mV to 0 mV. In these experiments, we applied 1 μM TTX, 20 mM TEA, 200 μM Cd^2+^ and 2.5 mM Cs^+^ to block Na_v_, most of K^+^, Ca_v_ and HCN channels, respectively. Subtracting the current obtained in XG from control current yielded the XG-sensitive current. We found no such current at negative potentials (n = 5, fig. 6). However, we did observe a *I_XG_*, from around −55 mV to less negative potentials (fig. 6c). This voltage range coincides with the ISI (Khaliq and Bean, 2008), suggesting that this current could correspond to the pacemaker current. Thus, the shape of *I_XG_* is inconsistent with it being an inward *I_ω_-*like current.

**Figure 6.**
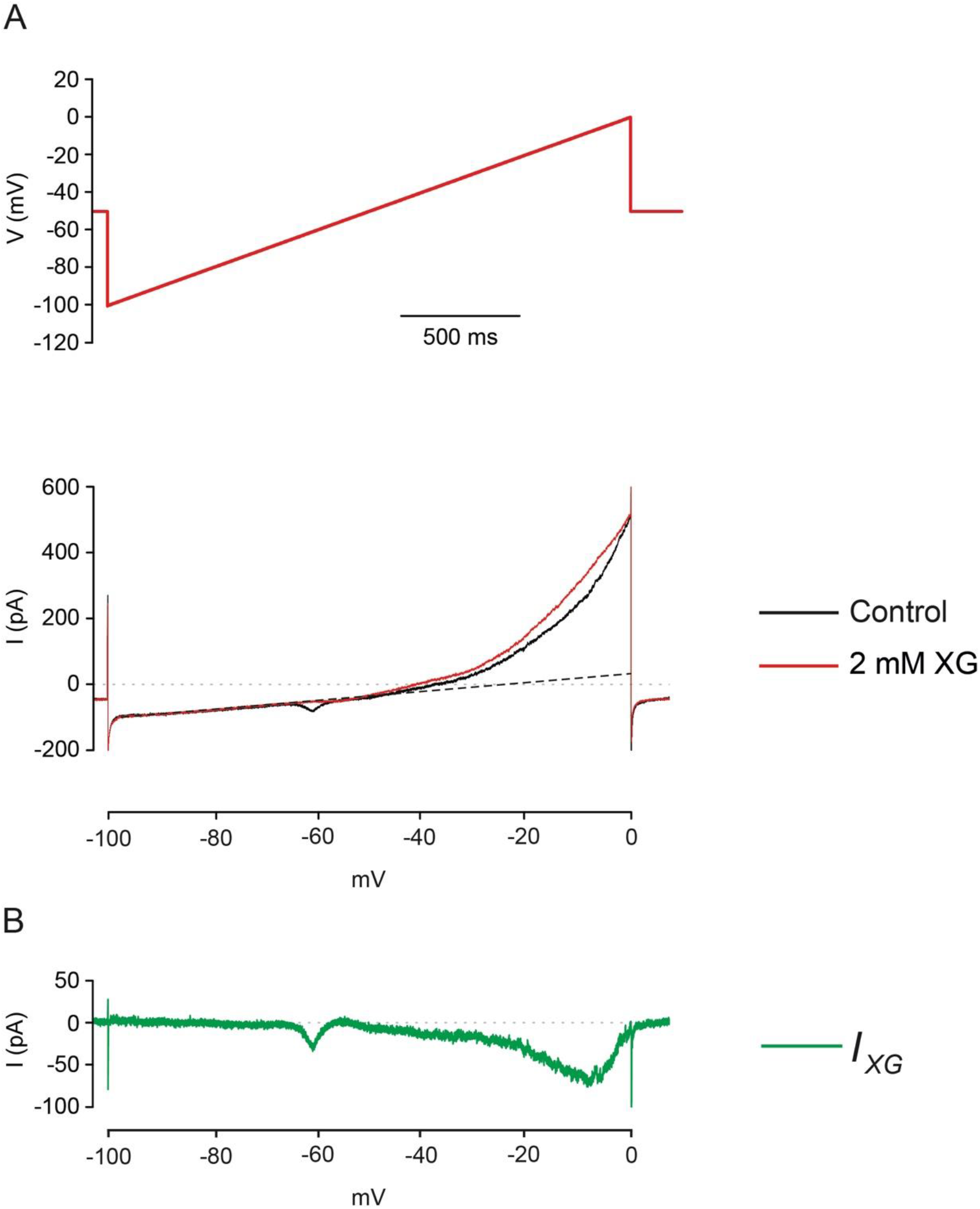
Isolation of I_XG_. **A.** Mean currents before (black) and after (red) application of 2 mM XG evoked in voltage-clamp by a ramp at a speed similar to the ISI (25 mV/s). In both conditions, 1 μM TTX, 20 mM TEA, 200 μM Cd^2+^ and 2.5 Cs^+^ were applied to block respectively Na_v_, most K^+^, Ca_v_ and HCN channels. Dotted line corresponds to the extrapolated linear regression between −98 mV and −70 mV. Its slope corresponds to an input resistance of 935 MΩ. The inward current at −60 mV corresponds to a T-type calcium current in control which is absent in the presence of XG, probably due to rundown. **B.** Subtraction of the control trace by the trace in the presence of XG in **A** reveals *I_XG_* plotted as a function of voltage. Note that the the current starts to operate at −50 mV.

Second, we addressed the possibility that the mRNA coding for an ion channel species could be edited physiologically by the type-2 adenosine deaminase acting on RNA (ADAR-2) enzyme. Such editing could convert an arginine of a S4 segment into a glycine. Indeed, ADAR-2 converts an adenosine into an inosine, which will be read as a guanosine. ADAR-2 is active in DA SNc neurons and has been shown to edit the IQ calmodulin-binding domain of Ca_v_ 1.3, modulating the inactivation of this channel by Ca^2+^ (Huang et al., 2012). We sequenced cDNA from SNc neurons and cortical neurons (as a negative control) in the region coding for the S4 segments of Na_v_1.1, 1.2 and 1.6 channels as there is ample evidence that pathological Na^+^ channels do generate *I_ω_* (Mason et al., 2019; Sokolov et al., 2007, 2005), and also sequenced the IQ calmodulin-binding domain of Cav 1.3 as a control. We found a small percentage of editing in some S4s, but this was observed in both SNc and cortex. On the other hand, we did find the heavy editing reported by Huang et al., demonstrating that our method was adequate (Tables 2 and S1). The fact that there was not a region-specific editing and the low percentage of editing indicate that ω pores are unlikely to be present in the tested channels. Taken together, our molecular and functional data are not in favor of the possibility that a typical gating pore current is the target of XG.

**Table 2.** Summary results from the sequencing of the various S4 segments of Na_v_1.1, 1.2 and 1.6 channels in SNc and cortex (n = 2). There is no evidence in favour of robust editing in these parts of the channels.

|  |  | 0 |  | 1 |  | 2 |  | 3 |  | 4 |  | 5 |  |
| --- | --- | --- | --- | --- | --- | --- | --- | --- | --- | --- | --- | --- | --- |
|  |  | SNc | Cortex | SNc | Cortex | SNc | Cortex | SNc | Cortex | SNc | Cortex | SNc | Cortex |
| SCN1A | S4.1 | - | - | R (100 %) | R (100 %) | R (100 %) | R (100 %) | R (100 %) | R (100 %) | K (100 %) | K (100 %) | - | - |
|  | S4.2 | - | - | R (100 %) | R (100 %) | R (100 %) | R (100 %) | R (100 %) | R (100 %) | K (100 %) | K (100 %) | K (100 %) | K (100 %) |
|  | S4.3 | K (100 %) | K (100 %) | R (100 %) | R (100 %) | R (100 %) | R (100 %) | R (100 %) | R (100 %) | R (100 %) | R (100 %) | - | - |
|  | S4.4 | R (100 %) | R (100 %) | R (100 %) | R (100 %) | R (100 %) | R (100 %) | R (100 %) | R (100 %) | R (100 %) | R (100 %) | K (100 %) | K (100 %) |
| SCN2A | S4.1 | - | - | R (100 %) | R (100 %) | R (100 %) | R (100 %) | R (100 %) | R (100 %) | K (100 %) | K (100 %) | - | - |
|  | S4.2 | - | - | R (100 %) | R (100 %) | R (100 %) | R (100 %) | R (100 %) | R (100 %) | K (100 %) | K (100 %) | K (100 %) | K (100 %) |
|  | S4.3 | K (100 %) | K (100 %) | R (100 %) | R (100 %) | R (100 %) | R (100 %) | R (98 %) / G (2 %) | R (99 %) / G (1 %) | R (100 %) | R (100 %) | - | - |
|  | S4.4 | R (100 %) | R (100 %) | R (100 %) | R (100 %) | R (100 %) | R (100 %) | R (100 %) | R (100 %) | R (100 %) | R (100 %) | K (100 %) | K (100 %) |
| SCN8A | S4.1 | - | - | R (100 %) | R (100 %) | R (100 %) | R (100 %) | R (100 %) | R (100 %) | K (100 %) | K (100 %) | - | - |
|  | S4.2 | - | - | R (100 %) | R (100 %) | R (100 %) | R (100 %) | R (100 %) | R (100 %) | K (100 %) | K (100 %) | K (100 %) | K (100 %) |
|  | S4.3 | K (100 %) | K (100 %) | R (100 %) | R (100 %) | R (100 %) | R (100 %) | R (100 %) | R (100 %) | R (100 %) | R (100 %) | - | - |
|  | S4.4 | R (100 %) | R (100 %) | R (100 %) | R (100 %) | R (100 %) | R (100 %) | R (100 %) | R (100 %) | R (100 %) | R (100 %) | K (100 %) | K (100 %) |

### XG blocks a current carried by Na^+^ and Cl^-^

Having identified *I_XG_* in DA SNc neurons in the relevant subthreshold range, our next aim was to define the permeant ions. To do so, while also attempting to mimic as closely as possible physiological conditions, we turned to a “pacemaker clamp” protocol (Philippart et al., 2016; Puopolo et al., 2007). In this voltage-clamp approach, the waveform of a spontaneously firing DA neuron is imposed as the command potential and the resulting currents are recorded. We then ran the voltage-clamp protocol in control solution and 2 mM XG and subtracted the current traces. These experiments were performed in TTX, TEA and Cd^2^. The internal control solution was K-gluconate-based (see methods). Consistent with our previous results, we observed a small inward *I_XG_* starting a few mV (−50 to −45 mV) below the threshold of the AP (fig. 7a). In control conditions, the amplitude of this current was – 23.85 ± 6.14 pA (n = 6) just below the onset of the action potential (−30 mV).

**Figure 7.**
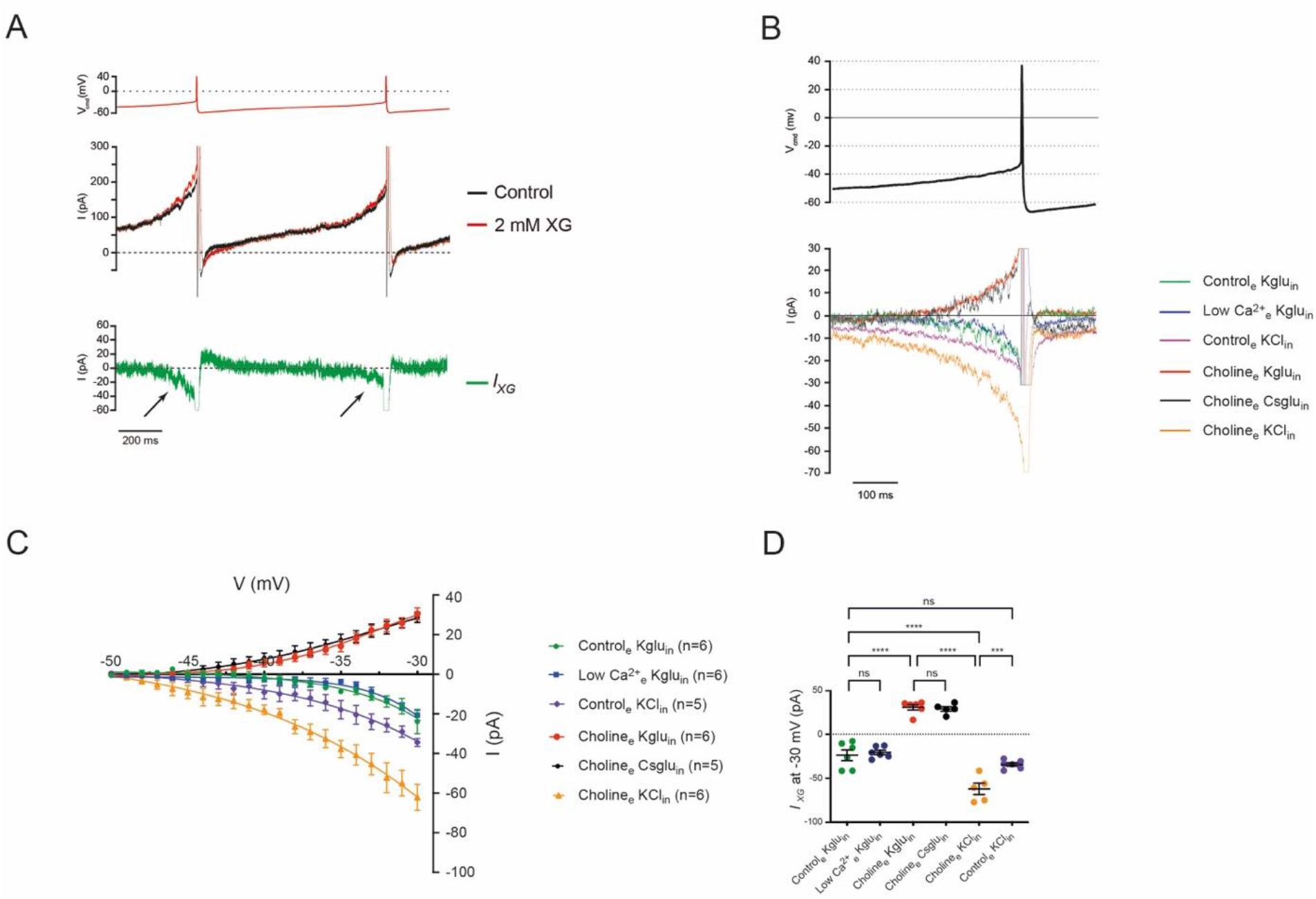
I_XG_ is activated during the ISI and is carried by Na^+^ and Cl^-^. **A.** Representative currents evoked by a pacemaking waveform in 1 μM TTX, 20 mM TEA and 200 μM Cd^2+^ before (black) and after (red) application of 2 mM XG (middle panel). Each trace represents an average of five sweeps obtained from one neuron. The subtraction of the control trace by the XG trace reveals *I_XG_.* Note that the latter starts below the AP threshold (arrows). **B.** Representative *I_XG_* evoked by a pacemaking waveform under different ion substitutions reveals that it is permeable to Na^+^ and Cl^-^. **C.** Summary results of *I_XG_* as a function of voltage and various ionic conditions. **D.** Summary results of *I_XG_* at −30 mV.

Next, we decreased the concentration of external Ca^2+^ ions from 2 to 0.1 mM. This procedure did not significantly change the amplitude of *I_XG_* (−20.73 ± 2.62 pA, n = 6, p = 0.83 versus control, ANOVA-1; fig. 7b-d), suggesting that *I_XG_* is not carried by Ca^2+^. We next substituted external Na^+^ by the much larger cation choline. This made *I_XG_* strongly outward (30.55 ± 3.01 pA in choline_e_ Kglu_in_ vs control_e_ Kglu_in_ at −30 mV, n = 6, p < 0.0001, ANOVA-1; fig. 7b-d). These results indicate that *I_XG_* is partially carried by Na^+^, in addition to another ionic species that produces the outward component.

Given the experimental conditions, K^+^ and Cl^-^ were the most plausible candidates. First, we substituted internal K^+^ by Cs^+^ in the external choline condition, in order to isolate the outward component. This did not change the outward component of *I_XG_* (28.80 ± 2.57 pA in choline_e_ Csglu_in_, n= 5, vs choline_e_ Kglu_in_, p = 0.83, ANOVA-1; fig. 7b-d), indicating that K^+^ ions probably do not contribute to this current. Next, we increased the internal Cl^-^ concentration from 25 mM to 145 mM resulting in a predicted shift of its reversal potential from ~ – 45 mV to 0 mV. Under these ionic conditions and in the presence of external choline, *IXG* was recorded as an inward current that was actually larger compared to the control condition (−62.10 ± 6.50 pA in choline_e_ KCl_in_, n = 6 vs control_e_ Kglu_in_, p <0.0001, ANOVA-1; fig. 7b-d). Taken together, these experiments demonstrated that *I_XG_* is carried by both Cl^-^ and Na^+^ ions. Finally, we recorded *I_XG_* in external Na^+^ and high internal Cl^-^. We hypothesized that, given that the two ionic species carry an inward current in these conditions, the global inward current would be larger than in the previous condition where only Cl^-^ could carry it. However, surprisingly, we observed a significantly smaller inward current compared to the “Cl^-^ only” condition (−34.40 ± 2.02 pA in control_e_ KCl_in_, n = 6, vs choline_e_ KCl_in_, p < 0.001, ANOVA-1, fig. 7b-d). On the other hand, the current amplitude was not significantly different from that of the control condition (controle KCl_in_ vs control_e_ Kglu_in_, p = 0.21, ANOVA-1; fig. 7b-d). These results strongly suggest that there is a competition between Na^+^ and Cl^-^ ions for the XG-sensitive pore. Therefore, Na^+^ and Cl^-^ ions generate a pacemaker current through one and the same pore.

### XG does not interact with the dopamine transporter

Given the results of our ion substitution experiments (fig. 7), we next turned to possible proteins that could sustain a low conductance permeable to both Na^+^ and Cl^-^, and that would be expressed only in DA neurons. The dopamine transporter (DAT) was an obvious candidate since it is only expressed in DA neurons. While transporting DA, it takes up Na^+^ and Cl^-^ in a 2:1 ratio, generating a small inward current that is able to modulate DA neuron excitability (Carvelli et al., 2004; Ingram et al., 2002; Sonders et al., 1997). In order to test the possible contribution of the DAT in *IXG*, we took two approaches.

First, we performed extracellular recordings in rat slices and applied 10 μM GBR12909, a specific blocker of DA uptake. The compound produced no significant change in firing frequency when the synaptic blockers (including sulpiride) were applied (2.77 ± 0.31 Hz in control vs 2.21 ± 0.23 Hz in 10 μM GBR12909, n = 6 for both conditions, p = 0.19, t-test; fig. 8a). Additionally, superfusion of 2 mM XG inhibited the firing of DA neurons to the same extent as in control conditions (from 2.28 ± 0.18 Hz in 10 μM GBR12909 to 0.01 ± 0.01 Hz in 2 mM XG, n = 6 for both conditions, p = 0.03, Wilcoxon test; fig. 8b). These observations suggest that uptake of DA is not generating the spontaneous firing and that DAT is probably not the target of XG.

**Figure 8.**
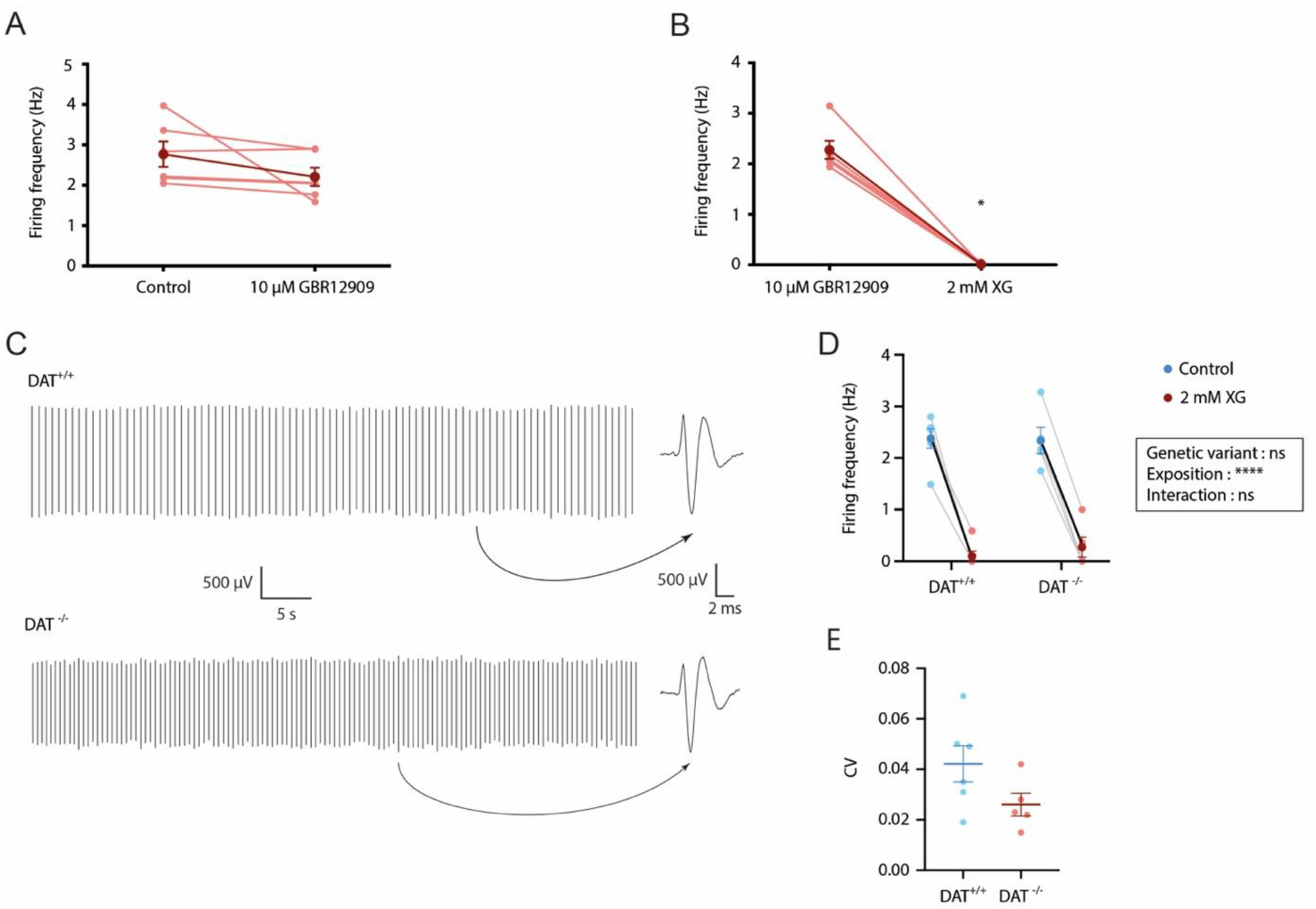
XG does not interact with the dopamine transporter. **A.** Summary data of the effect of 10 μM GBR12909 on the firing frequency of DA SNc neurons. The individual frequencies are represented in light red (n = 6 for both conditions) while the means ± SEM are represented in dark red. **B.** Summary data of the effect of 2 mM XG on the firing frequency of DA SNc neurons during application of 10 μM GBR12909. The individual frequencies are represented in light red (n = 6 for both conditions) while the means ± SEM are represented in dark red. **C.** Representative extracellular recordings of DA neurons from DAT^+/+^ (upper panel) and DAT^-/-^ (down panel) mice. Insets show the waveform of an action potential. **D.** Summary data of the effect of 2 mM XG on the firing frequency of DA SNc neurons from DAT^+/+^ and DAT^-/-^ mice. The individual frequencies are represented in light color (n = 6 for both DAT^+/+^ and DAT^-/-^ mice) while the means ± SEM are represented in dark color. **E.** Summary data of the coefficient of variation (CV) of the firing frequency of DA SNc neurons from DAT^+/+^ and DAT^-/-^ mice. The individual frequencies are represented in light color (n = 6 for both DAT^+/+^ and DAT^-/-^ mice) while the means ± SEM are represented in dark color.

Second, and in order to confirm those results, we performed extracellular recordings on wild type and DAT^-/-^ mice. DA neurons from DAT^-/-^ mice were spontaneously active (fig. 8c). Their firing rate was similar to the one of WT neurons (2.38 ± 0.19 Hz for DAT^+/+^, n = 6 vs 2.34 ± 0.26 Hz for DAT^-/-^, n = 5; fig. 8c,d), as was their regularity (coefficient of variation of the ISI [see methods] was 0.042 ± 0.007 for DAT^+/+^, n = 6 vs 0.026 ± 0.005 for DAT^-/-^, n = 5, p = 0.1, t-test; fig. 8e). In addition, XG inhibited the firing of DA neurons from the two genotypes to a similar extent (p = 0.52 for genetic variant, ANOVA-2; fig. 8d). Taken together, these two types of experiments demonstrate that XG does not interact with the DAT and that the latter is not responsible for the spontaneous activity of DA SNc neurons.

## DISCUSSION

We have tested the hypothesis that a compound previously reported to be a gating pore blocker, XG, is able to inhibit the spontaneous slow pacemaking of DA SNc neurons. Our results show that this is indeed the case in both rat and mouse DA neurons and in slices taken from both juvenile and adult animals. The IC_50_ of the compound is in the high micromolar range and consistent with the pharmacology of *I_ω_* (Sokolov et al., 2010). Using two different salts of XG allowed us to establish that it is the xylylguanidinium moiety that is responsible for the observed inhibition. Moreover, we found that XG inhibits pacemaking of DA neurons without hyperpolarizing them and without increasing their whole-cell conductance (contrary to what a K^+^ channel opener would do). Finally, we found that XG displays a reasonable degree of specificity, since fast GABAergic pacemaker neurons of the SNr were unaffected by the drug. Therefore, we may have found the first specific inhibitor of the spontaneous activity of DA neurons. In addition, XG inhibits pacemaking of DA neurons, but does not affect their bursting behavior. To our knowledge, this is the first compound that is able to do so. In addition, the XG-sensitive current has a shape that is fundamentally different from what was previously reported for inward gating pore currents (Sokolov et al., 2010). An effect of XG via inhibition of NALCN channels could also be excluded. Intriguingly, we demonstrated that the XG-sensitive current is carried by both Na^+^ and Cl^-^. However, although the DAT transports Na^+^ and Cl^-^ during dopamine uptake, we could rule out that this protein sustains slow pacemaking in DA neurons.

Mesencephalic DA neurons constitute a heterogeneous population in terms of electrophysiological and molecular properties. The electrophysiological characteristics of mesencephalic DA neurons, including AP duration and the density of *I_A_* and *I_H_,* vary along a medio-lateral axis going from mesocortical projections (medial) to nigrostriatal projections (lateral) (Amendola et al., 2012; Lammel et al., 2008; Margolis et al., 2008; Neuhoff et al., 2002; Tarfa et al., 2017). In addition, mesoprefrontal VTA DA neurons in mice (Lammel et al., 2008) and amygdala-projecting VTA DA neurons in rats (Margolis et al., 2008) display unique properties, such as lack of somato-dendritic D2 receptors and high-frequency pacemaking. As far as the slow pacemaking activity is concerned, various types of voltage-dependent channels have been suggested or shown to contribute to it, including several types of Ca_v_ channels, e.g. L-type channels in some cells and P/Q type channels in other cells (Puopolo et al., 2007), as well as HCN channels (Neuhoff et al., 2002; Seutin et al., 2001). In addition, the ionic nature of the channels operating during the ISI differs between VTA and SNc DA neurons (Khaliq and Bean, 2010; Puopolo et al., 2007), even though the spontaneous activity is remarkably similar in DA neurons of both regions. In this study, we found a complete or near complete inhibition of firing with 2 mM XG in all neurons that we studied. We suggest that the XG-sensitive conductance provides the depolarizing drive in all slow DA pacemakers. Thus, these neurons, regardless of their location within the midbrain, would be endowed with a robust common pacemaking mechanism, on top of which additional conductances would be expressed in a region-specific manner.

XG has only been described so far as a blocker of *I_ω_* (Sokolov et al., 2010). Regarding the effect of *I_ω_* on cell excitability, it has been shown that R222W in hNav1.4 channels shifts the resting potential of muscle fibers to more depolarized state, facilitating the generation of APs (Bayless-Edwards et al., 2018). The amplitude of APs is also decreased as the sustained inward current reduces Nav channel availability. In the brain, R853Q hNav1.2 channels have been shown to generate a K^+^-selective inward current linked to epilepsy (Mason et al., 2019; Sokolov et al., 2005). Our data show no evidence for an *Iω*, as we did not observe XG-sensitive inward currents at hyperpolarized potential (fig. 6c) and the sequencing of Nav channels S4 did not show any R/X variant (Table 2). However, we cannot exclude that editing occurs in DA neurons in channels that were not investigated in our study, as ω pores were observed previously in shaker K^+^ channels (Starace and Bezanilla, 2004; Tombola et al., 2005). Note that a combined permeability to cations and anions has been previously demonstrated for *I_ω_* (Khalili-Araghi et al., 2012). Therefore, our results do not formally exclude that a gating pore current underlies pacemaking in DA neurons. Other possibilities exist. For example, it has been shown that the biophysical properties of voltage-gated proton channels are similar to the ones of ω pores (Hong et al., 2013). Since they are blocked by guanidinium derivatives, the structures of which are very similar to the one of XG, it is possible that such channels are the target of XG.

Even though we did not identify the molecular target of XG, we characterized its ion permeability and found that both Na^+^ and Cl^-^ permeate through the same pore. This mechanism may be critical to ensure the temporal stability of pacemaking. Thus, any depolarizing event will increase the ratio of Cl^-^ versus Na^+^ influx and the converse will be true for a hyperpolarizing event. This is what we would expect from a pacemaker current in order to sustain a stable spontaneous activity. In light of our results, we speculate that *IXG* might correspond to the small inward current in the ISI described by others (Khaliq and Bean, 2008). Future application of XG to other slow pacemaker cells will clarify if the mechanism proposed here is of more general relevance. Additionally, we expect that a full characterization of *IXG*, when identified, can be used to verify the mechanism we propose in a computational model of DA neuron pacemaking.

In conclusion, we provide evidence for the involvement of an unconventional conductance in the generation of slow pacemaking of DA neurons. This conductance is sensitive to a blocker of *I_ω_*, yet lacks some characteristics of gating pores, and is permeable to both Na^+^ and Cl^-^, a feature that, to our knowledge, has not been reported so far for a membrane ion channel.

## MATERIALS AND METHODS

### Animals

All our experiments were performed in accordance with the ethical guidelines of the European Union (Directive 2010/63/EU). In addition, all procedures of animal care, handling and sacrifice were approved by the Ethics Committee for Animal Use of Liège University (protocol 1210)

Both juvenile (18-26 days) and adult (6-8 weeks) Wistar rats were used for patchclamp and extracellular recordings, respectively. DA neurons reach their electrophysiological maturity at P18 (Dufour et al., 2014; Tepper et al., 1994). Juvenile C57Bl6/N mice were used as well for both types of recordings. A mouse model bearing a complete KO for the DAT was also used in extracellular recordings. This mouse line (Giros et al., 1996) was kindly provided by Pr. Marc Caron (Duke University, Durham, NC, USA) and has been amply characterized.

### Molecular biology

Parts of the SNc and cortex were isolated from adult rat brains and kept at −80°C. RNA was extracted using the “Monarch Total RNA Miniprep” kit (New England Biolabs, Ipswich, MA, USA) and then was reverse transcribed by random priming using the PrimeScript RT Reagent Kit (Takara Bio Europe, Saint-Germain-en-Laye, France). PCRs were performed using the Q5 polymerase (New England Biolabs) and specific primer pairs (Integrated DNA Technologies, Leuven, Belgium). Specificity of amplification was first assessed by gel agarose analysis and then PCR fragments were processed by the GIGA-Genomics platform (GIGA Institute, Liège University) for high throughput sequencing on an Illumina sequencer.

The sequences of primer pairs that were designed to amplify the fourth alpha-helices of each transmembrane domain of the voltage-gated Na^+^ channels and of the HCN channels, or to amplify the IQ domain of the voltage-gated Ca^2+^ channel, are given in Table S1. Delineations of alpha-helices or of the IQ domain are given under the accession number describing each channel.

### Electrophysiology

Animals were anaesthetized with isoflurane and sacrificed by decapitation. The brain was quickly removed and submerged in an oxygenated ice-cold physiological saline solution (ACSF) containing the following (in mM): 125 NaCl, 2.5 KCl, 1 MgCl_2_, 2 CaCl_2_, 10 (extracellular recordings) or 25 (patch clamp recordings) glucose, 1.25 NaH_2_PO_4_, 25 NaHCO_3_, pH 7.4 (95% O_2_ 5% CO_2_ and 310 mosm/L). Coronal or horizontal 300 μM-thick slices were obtained using a vibratome (Leica VT-1200, Nussloch, Germany) for patch-clamp and extracellular recordings respectively. Midbrain slices were incubated at 34°C for 30-60 min before being stored at room temperature.

Extracellular recordings were performed as previously described (Drion et al., 2011). DA neurons were identified based on their low firing rate (0.5-5 Hz), long AP duration (> 2 ms) and very regular pacemaking activity (CV_ISI_ < 10 % over a 5 min period). Numerous previous experiments have shown that these neurons are inhibited by dopaminergic agonists via D2 receptors and are therefore considered as DA. Neurons whose firing frequency varied by more than 5 % in control conditions were discarded. To quantify the effect of various concentrations of XG on the firing frequency, we calculated the mean frequency of the neurons over a 5-min control period. We next measured the mean frequency of the neuron during the last minute of superfusion of a given drug concentration and calculated the ration of the two values as a percentage. Due to the difficulty of solubilizing XG-carbonate, each neuron was exposed to only one concentration of that salt. Briefly, the ACSF had to be brought to a pH of ~3 for a full dissolution to occur. The pH was then brought back to 7.4 while the ACSF was oxygenated with carbogen and heated. In order to exclude artifacts related to pH manipulations, a control solution that did not contain the drug was prepared in parallel and superfused as the control solution. On the other hand, XG-mesylate solubilized easily at physiological pH. Therefore, each neuron was exposed to various concentrations of this salt. To quantify firing regularity in the DAT KO experiments, we measured the CVISI defined as followed: CVISI = SDISI/meanISI. CVISI was measured during the control period for 5 min, before superfusion of XG.

For whole-cell patch-clamp recordings, slices were positioned in the recording chamber and superfused with heated ACSF (34°C). Neurons were visualized using infrared-Dodt gradient contrast (IR-DGC) optics on a Zeiss Axio Examiner A1 microscope equipped with a CCD camera (C7500-51; Hammamatsu, Japan). Patch pipettes (3-5 MΩ) were pulled from filamented borosilicate glass tubing (2.0 mm outer diameter, 0.42 mm wall thickness; Reference 1403516, Hilgenberg, Germany) with a horizontal puller (P-97; Sutter Instruments, Novato, CA, USA). Pipettes were filled with a potassium gluconate-based solution, consisting of (in mM): 125 K-gluconate, 20 KCl, 10 HEPES, 4 Mg-ATP, 0.3 Na-GTP, 10 Na2-phosphocreatine, and 0.5 EGTA, pH 7.2 (~ 300 mosm/L). Patch-clamp recordings were performed using a Multiclamp 700B amplifier connected to a PC through a Digidata 1550 interface (Molecular Devices, San Jose, CA, USA) and we acquired data with pClamp 10.5 (Molecular Devices). Data were sampled at 50 kHz and filtered at 10 kHz through a Bessel filter. Some traces in the figures were digitally filtered at 2 kHz (Gaussian characteristics). Analyses were carried out in pClamp 10.5 (clampFit) or Stimfit 0.15 (Christoph Schmidt-Hieber, UCL).

SNc DA neurons were identified based on their electrophysiological properties, including low frequency of firing (0.5-5 Hz), AP duration (width at half-amplitude > 1.35 ms) and the presence of a strong hyperpolarization-activated inward current (*IH*) (fig. S1) (Richards et al., 1997; Seutin and Engel, 2010). Burst recordings were performed as previously described (Destreel et al., 2019; Johnson et al., 1992).

Intracellular application of XG mesylate was done by adding it to the intracellular solution. For these experiments, a neuron was first patched in the whole-cell configuration with the control intracellular solution to check its DA nature (see above) and record its basal spontaneous activity. Then, the pipette was withdrawn from the neuron and the neuron was patched with a second pipette containing the drug.

For pacemaker clamp recordings, patch pipettes had a very low resistance (1-3 MΩ) and various solutions were used to determine the ionic nature of *IXG*. For the external solution, we first used the standard ACSF solution (see above). The low Ca^2+^ solution contained 0.1 mM CaCl_2_; for the high choline solution, we replaced NaCl equimolarly by choline chloride. For the internal solution, we used a potassium gluconate-based solution (as described above), a cesium gluconated-based solution, in which K-gluconate and KCl were replaced equimolarly by Cs-gluconate and CsCl, respectively and a potassium chloride-based solution in which K gluconate was replaced by KCl, giving a total intracellular concentration of 145 mM KCl. Neurons whose access resistance increased by >20 % over the duration of the recording were discarded. All the experiments described above, expect bursting activity recordings, were performed in the presence of synaptic blockers to isolate DA neurons pharmacologically and to prevent any presynaptic effect. We used 10 μM CNQX, 10 μM SR95531, 1 μM MK801, 1 μM sulpiride, and 1 μM CGP55845, to block respectively AMPA/kainate, GABAA, NMDA, D_2_/D_3_, and GABAB receptors.

For cell-attached recordings, 250 μm-thick coronal midbrain slices were obtained from 10 to 12 week-old male C57Bl6N mice. Animals were sectioned in ice-cold ACSF (containing in mM: 50 sucrose, 125 NaCl, 2.5 KCl, 25 NaHCO_3_, 1.25 NaH_2_PO_4_, 2.5 glucose, 6.2 MgCl_2_, 0.1 CaCl_2_, and 3 kynurenic acid; Sigma-Aldrich GmbH; bubbled with 95% 0_2_ / 5% CO_2_) after intracardial perfusion with ice-cold ASCF. After sectioning, slices were rested for recovery at least one hour before recording in 95% 02 /5% CO_2_ ACSF at 37°C. Recordings were carried out in ACSF (containing in mM: 22.5 sucrose, 125 NaCl, 2.5 KCl, 25 NaHCO_3_, 1.25 NaH_2_PO_4_, 2.5 glucose, 2 MgCl_2_, 2 CaCl_2_; bubbled with 95% 0_2_ / 5% CO_2_ in the presence of CNQX (20 μM), DL-AP-5 (4 μM) and Gabazine (SR95531, 4 μM) to inhibit fast excitatory and inhibitory transmission. DA SNc neurons were initially visually identified and recorded in cell-attached mode and subsequently in whole-cell using a pipette solution containing in mM: 135 K-gluconate, 10 HEPES, 5 KCl, 5 MgCl_2_, 0.075 CaCl_2_, 0.1 EGTA, 5 ATP, 1 GTP, 0.1% neurobiotin.

Human NALCN, UNC79, UNC80, FAM155A complementary DNAs (cDNAs), codon optimized for *Homo sapiens* in pCDNA3.1(+) vectors, were used as previously described (Chua et al., 2020). The NALCN, UNC-79, UNC-80, and FAM155A RNAs were mixed in a ratio of 1:1:1:1 and injected into *Xenopus laevis* oocytes. Injected cells were incubated in ND96 (96 mM NaCl, 2 mM KCl, 1 mM MgCl_2_, 1.8 mM CaCl_2_, and 5 mM HEPES; pH 7.4) supplemented with 0.5 mM theophylline, 2.5 mM Na pyruvate, 50 μg/mL gentamycin and 50 μg/mL tetracycline at 18 °C, 140 rpm. Two-electrode voltage-clamp measurements were performed on oocytes five days after injection using a Warner OC-725C Oocyte Clamp amplifier (Warner Instrument Corp, USA). Oocytes were continuously perfused in a Ca^2+^ /Mg^2+^ -free ND96 recording solution (96 mM NaCl, 2 mM KCl, 2 mM BaCl_2_, and 5 mM HEPES; pH 7.4) at room temperature. Data were acquired using the pCLAMP 10 software (Molecular Devices) and a Digidata 1550 digitizer (Molecular devices), sampled at 10 kHz. Electrical powerline interference was filtered with a Hum Bug 50/60 Hz Noise Eliminator (Quest Scientific). Recording microelectrodes with resistances around 0.2 to 1.0 MΩ were pulled from borosilicate glass capillaries (Harvard Apparatus) using a P-1000 Flaming/Brown Micropipette Puller System (Sutter Instrument) and were filled with 3 M KCl. Stock solutions (1 M) of gadolinium (Gd^3+^, Sigma Aldrich) and XG-mesylate were prepared in Ca^2+^ /Mg^2+^ -free ND96 and diluted to desired concentrations.

### Immunofluorescence & confocal microscopy

Cell-attached recordings were converted to the whole-cell configuration to fill neurons with 0,1% neurobiotin for post-hoc verification of anatomical and neurochemical identity. The slices were then fixed in 4% PFA-PBS solution (containing 15% Picric acid, 0.1 M buffer, pH = 7.4) over night at 4°C, rinsed in PBS (containing in mM: 137 NaCl; 2.7 KCl; 10 NaH_2_PO_4_; 10 Na_2_HPO_4_; pH = 7.4) and incubated in blocking solution (0.2 M PBS with 10% horse serum, 0.5% Triton X-100, 0.2% BSA) for 1 hour. Slices were incubated with polyclonal anti-tyrosine hydroxylase rabbit antibodies (1:1000, Calbiochem, Merck) and were incubated overnight in carrier solution (containing in PBS: 0.2% PSA, 0.5% Triton, 1% horse serum) at room temperature. Slices were again rinsed in PBS and incubated in carrier solution with AlexaFluor-568 secondary anti-rabbit polyclonal goat antibody (1:1000; Thermo Fisher Scientific) and Streptavidin AlexaFluor-488 (1:750; Thermo Fisher Scientific) over night, then rinsed and mounted in Vectashield. Confocal microscopy was used to identify recorded neurons as dopaminergic and locate the position of neurobiotin-positive recorded neurons inside the SNc.

### Synthesis of XG-mesylate

#### N,N’-diBOC-2,4-Xylyl Guanidine (BOC-XG) preparation

2,4-dimethylaniline (about 50 mmol) reacted overnight with N,N’-diBOC-1H-pyrazole-1-carboxamidine (1.2 equivalents) as well as N-ethyldiisopropylamine (2 equivalents) in 100ml of methanol. The reaction medium was then evaporated under reduced pressure. The resulting crude oil was dissolved in 100 ml ethyl acetate and washed three times by 30 ml HCl (1 N) and by 50 ml brine. The organic solution was evaporated under reduced pressure, and the residue was then crystallized in methanol.

**^1^H NMR** (CDCl_3_,500 MHz) □ (ppm) 1.47 (s, 10H), 1.54 (s, 10H), 2.26 (s, 3H), 2.28 (s, 3H), 7.00 (m, 2H), 7.75 (d, J = 8.14 Hz, 1H), 10.04 (s, 1H), 11.67 (s, 1H) **^13^C NMR**: (CDCl_3_,500 MHz) □ (ppm) (CH_3_)18.13, 20.82, 28.10, 28.22, (CH)124.64, 127.06, 130.95, (C) 79.38, 83.42, 130.35, 132.49, 134.96, 153.44, 154.03, 163.78 **Elemental Analysis:** C_19_H_29_N_3_O_4_ (363.46 g/mol) Calculated (%) C 62.79, H 8.04, N 11.56, Found (%) C 62.58, H 8.28, N 11.63

#### 2,4-Xylyl Guanidine mesylate (XG mesylate) preparation

The isolated BOC-XG was dissolved in 50 ml dichloromethane and methanesulfonic acid (2.5 ml per gram) was added. After 4 to 5 hours at room temperature, the medium was evaporated under reduced pressure, the oily residue was subsequently washed five times by about 20 ml diisopropylether and dried under reduced pressure. The XG mesylate was then crystallized in hot ethanol by addition of cold diethylether.

**^1^H NMR**: (CDCl_3_,500 MHz) □ (ppm) 2.27 (s, 3H), 2.32 (s, 3H), 2.85 (s, 3H), 7.07 (m, 3H), 7.58 (broad, 3H), 9.28 (s, 1H)

**^13^C NMR**: (CDCl_3_,500 MHz) □ (ppm) (CH3) 17.18, 20.88, 39.22, (CH) 127.46, 128.00, 132.36, (C) 129.80, 135.96, 138.88, 156.93

**Melting point**: 105.5–107.5°C (several samples presented another polymorphic form, confirmed by Raman and FTIR spectra, and a melting point at 137.5-139.0°C)

**Elemental Analysis:** C_10_H_17_N_3_O_3_S (259.33 g/mol) Calculated (%) C 46.32, H 6.61, N 16.20, S 12.36 Found (%) C 46.33, H 6.70, N 16.25, S 12.02

**Mass Spectrometry**: C_9_H_13_N_3_ (163.11 g/mol) Adduct +H^+^, extraction mass 164.11822 Da, found at 164.11817 Da

### Statistics

Data are presented as mean ± standard error of the mean (SEM). Statistical analyzes were performed using Prism 8. Normality tests were performed prior to analyzing the data by either a parametric or non-parametric test. In this study, we used Student t-test/Wilcoxon test, ANOVA-1/Friedman followed by a post hoc Holm-Sidak’s/Dunn’s test and ANOVA-2 when required. Significance level is given as *<0.05, **<0.01, ***<0.001, ****<0.0001 and is displayed in the figures.

### Chemicals

Apamin, CNQX and SR9551 were purchased from Alomone labs (Jerusalem, Israel); isradipine, ZD 7228, MK801, NMDA, sulpiride and CGP55845 from Tocris Bioscience (Bristol, UK); TTX and Cs-gluconate from Hello Bio (Bristol, UK); 4-AP, nifedipine, XG carbonate and salts used to prepare the external and internal solutions were obtained from Sigma-Aldrich (Overijse, Belgium).

## DATA AVAILABILITY

Access to data files and statistical analysis will be provided upon full publication.

## ACKNOWLEDGEMENTS

We thank all the members from the laboratory of Neurophysiology for the helpful discussions and their suggestions. We are also grateful to the GIGA-Genomics platform for the sequencing of mRNA. This work was supported by grants from the “Fonds National de la Recherche Scientifique” (FNRS, Belgium) (J.0148.19 to VS VS), from the “Fondation Léon Frédéricq” (Belgium) (FHULF-D.MESGCAN.01-05 and “prix de l’espoir” to KJ). JFL is a Research Director of the FNRS. DE and BL are Research Associates of the FNRS. KJ, RV and SR received a grant from the “Fonds de la Recherche dans l’Industrie et l’Agriculture” (FRIA).

## EXTENDED DATA

**Figure S1.**
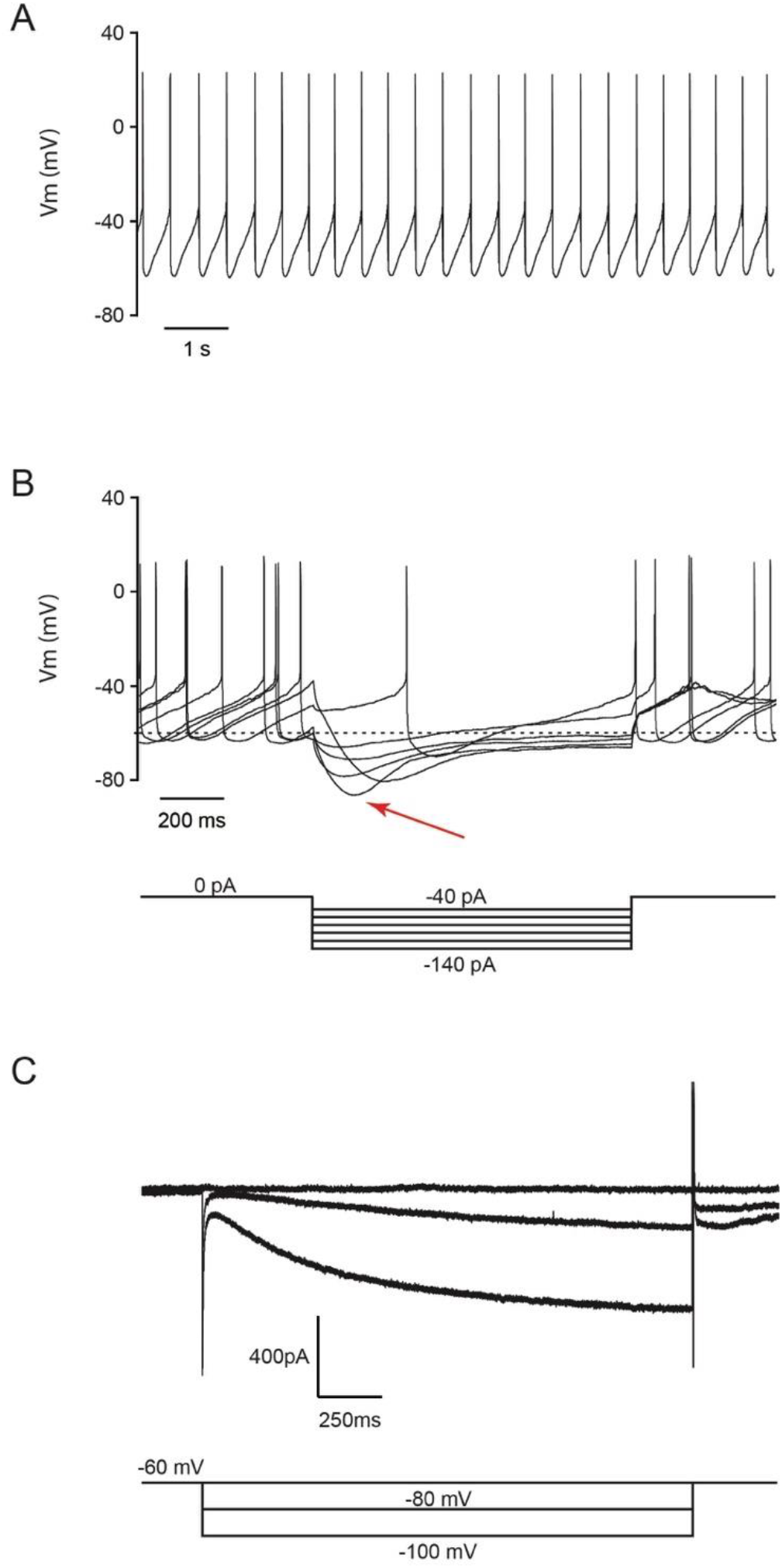
Electrophysiological identification of DA SNc neurons. **A.** Representative recording of spontaneous activity from a DA SNc neuron. **B.** Example of voltage traces evoked by a 1s current injection from −140 pA to −40 pA in 20 pA steps. The red arrow points to the membrane rectification, or *sag* due to the activation of HCN channels. **C.** Representative recording of *I_h_* evoked by a 2 s step from −60 mV to −100 mV (represented below). The slow pacemaker activity and the presence of *I_h_* support the DA identity of the recorded neurons (Destreel et al., 2019).

**Figure S2.**
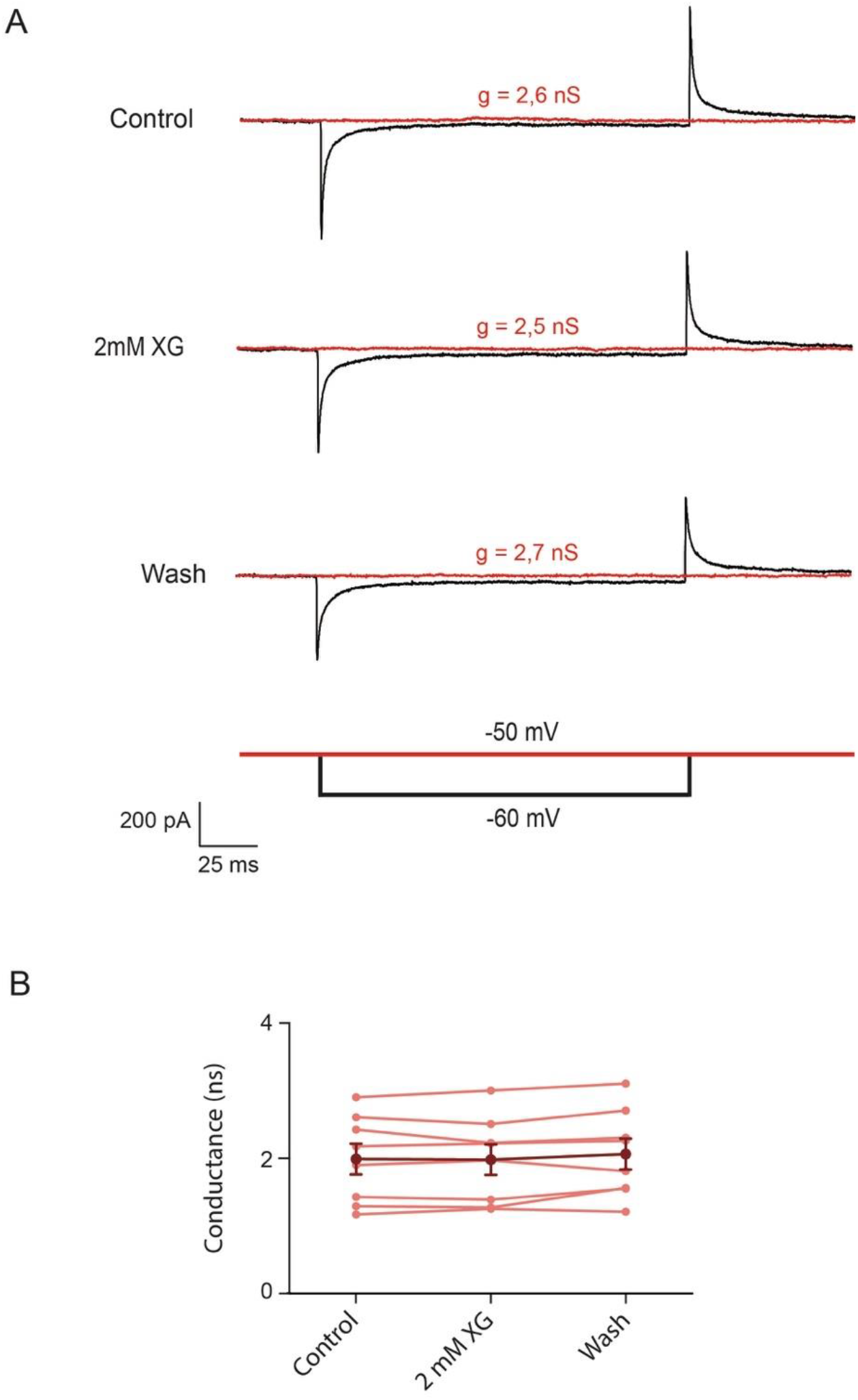
XG does not increase the whole-cell conductance of DA SNc neurons. **A.** Representative recording of currents evoked by a 100 ms step from −50 mV (red) to – 60 mV (black). The conductance calculated from g = *d*I/*d*V. **B.** Summary data of the effect of 2 mM XG on the conductance. The light red data corresponds to individual neurons (n = 8) and the dark red corresponds to the mean ± SEM of all recorded neurons.

**Figure S3.**
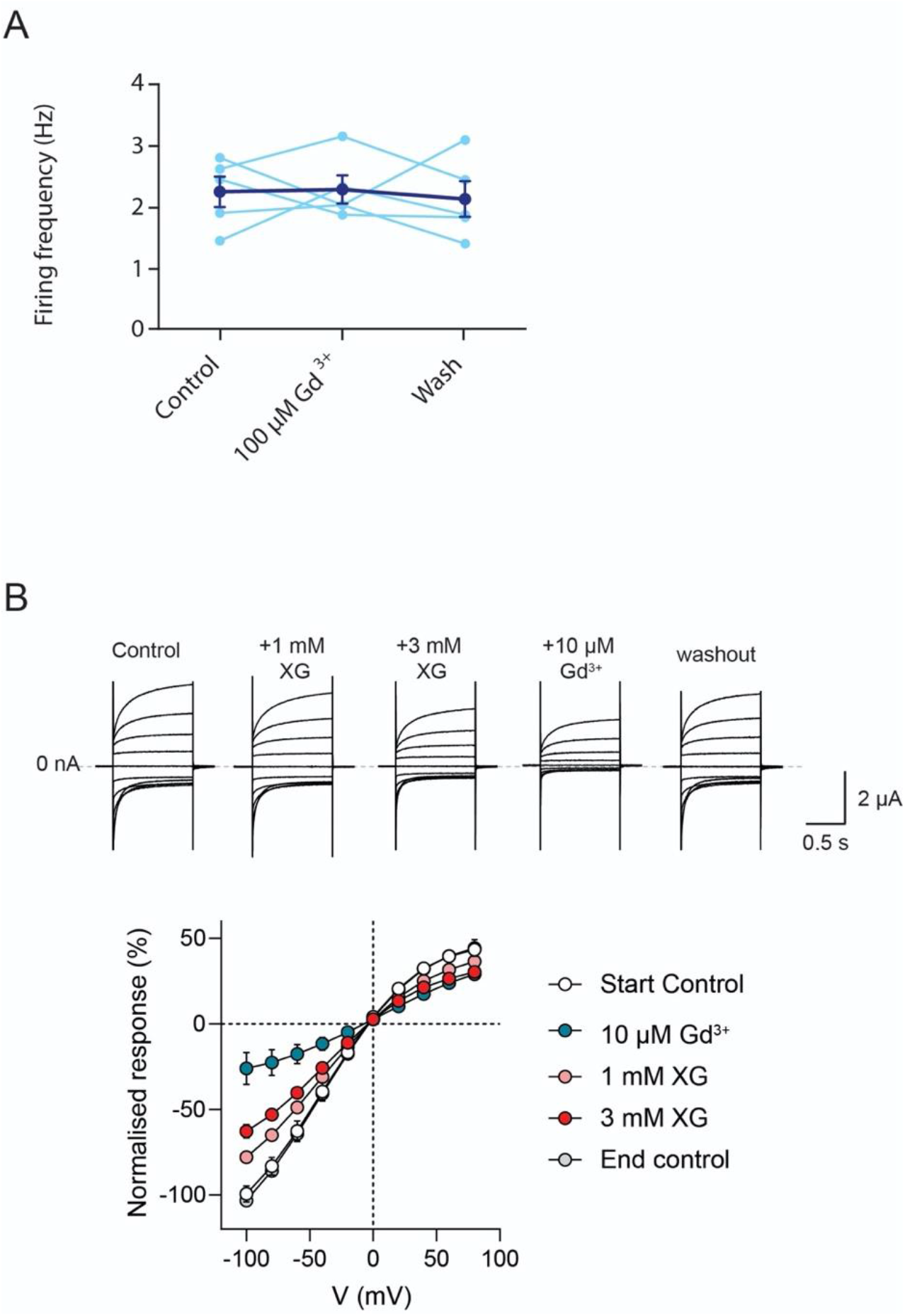
Inhibition of pacemaking of SNc neurons does not involve NALCN channels. **A.** Summary data of the effect 100 μM Gd^3+^ on the firing frequency of DA SNc neurons. The light blue data corresponds to individual neurons (n = 5) and the dark blue corresponds to the mean ± SEM of all recorded neurons. **B.** Representative traces of NALCN currents during application of XG and Gd^3+^ (top). Summary plot of the effects of XG and Gd^3+^ on NALCN-mediated instantaneous inward currents expressed in *Xenopus laevis* oocytes (bottom).

**Table S1.** Summary results from the sequencing of the IQ domain of Ca_v_1.3 channels in SNc and cortex (n = 2). These data are consistent with what has already been showed by others (Huang et al., 2012).

| Cav 1.3 IQ domain editing |  |  |  |
| --- | --- | --- | --- |
|  |  | SNc | Cortex |
| *CACNA1D*IQ | I / M | 68 % / 32 % | 81 % / 19 % |
|  | Q / R | 97 % / 3 % | 99 % / 1 % |
|  | Y / C | 78 % / 22 % | 87 % / 13 % |

**Table S2.**
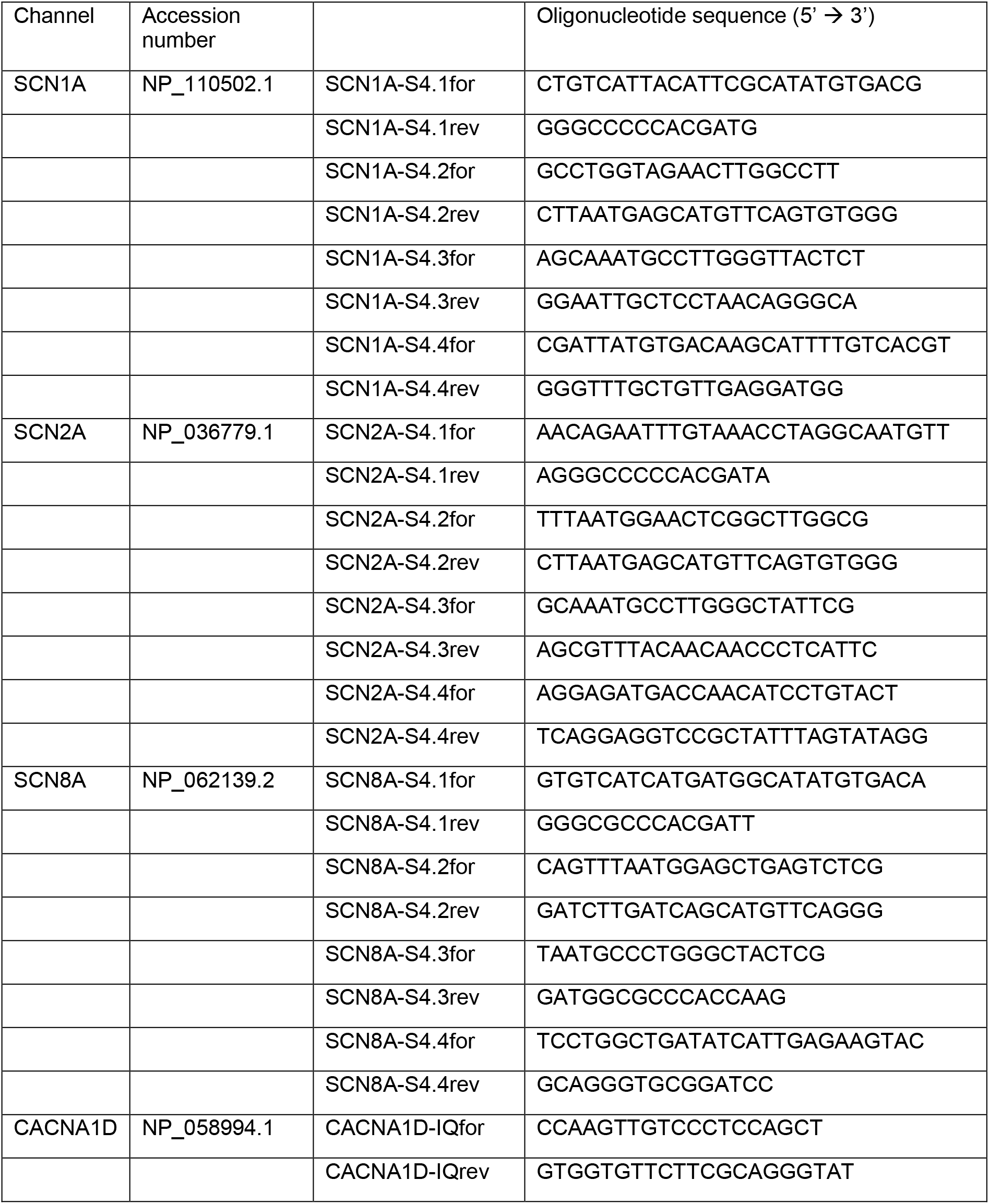
Sequences of primer pairs used to amplify the S4 transmembrane domains of various Na_v_ channels or the IQ domain of Ca_v_1.3 channels.

## REFERENCES

Albin RL, Young AB, Penney JB. 1989. The functional anatomy of basal ganglia disorders. Trends Neurosci 12:366–375. doi:10.1016/0166-2236(89)90074-X

Amendola J, Woodhouse A, Martin-Eauclaire MF, Goaillard JM. 2012. Ca 2+/cAMP-sensitive covariation of I A and I H voltage dependences tunes rebound firing in dopaminergic neurons. J Neurosci 32:2166–2181. doi:10.1523/JNEUROSCI.5297-11.2012

Bayless-Edwards L, Winston V, Lehmann-Horn F, Arinze P, Groome JR, Jurkat-Rott K. 2018. NaV1.4 DI-S4 periodic paralysis mutation R222W enhances inactivation and promotes leak current to attenuate action potentials and depolarize muscle fibers. Sci Rep 8:1–13. doi:10.1038/s41598-018-28594-5

Blythe SN, Atherton JF, Bevan MD. 2007. Synaptic Activation of Dendritic AMPA and NMDA Receptors Generates Transient High-Frequency Firing in Substantia Nigra Dopamine Neurons In Vitro. J Neurophysiol 97:2837–2850. doi:10.1152/jn.01157.2006

Boone AN, Senatore A, Chemin J, Monteil A, Spafford JD. 2014. Gd3+ and calcium sensitive, sodium leak currents are features of weak membrane-glass seals in patch clamp recordings. PLoS One 9. doi:10.1371/journal.pone.0098808

Carvelli L, Mcdonald PW, Blakely RD, Defelice LJ. 2004. Dopamine transporters depolarize neurons by a channel mechanism. Proc Natl Acad Sci 101:16046–16051. doi:10.1073/pnas.0403299101

Chan CS, Guzman JN, Ilijic E, Mercer JN, Rick C, Tkatch T, Meredith GE, Surmeier DJ. 2007. ‘Rejuvenation’ protects neurons in mouse models of Parkinson’s disease. Nature 447:1081–1086. doi:10.1038/nature05865

Chua HC, Wulf M, Weidling C, Rasmussen LP, Pless SA. 2020. The NALCN channel complex is voltage sensitive and directly modulated by extracellular calcium. Sci Adv 6. doi:10.1126/sciadv.aaz3154

de Vrind V, Scuvée-Moreau J, Drion G, Hmaied C, Philippart F, Engel D, Seutin V. 2016. Interactions between calcium channels and SK channels in midbrain dopamine neurons and their impact on pacemaker regularity: contrasting roles of N-and l-type channels. Eur J Pharmacol 788:274–279. doi:10.1016/j.ejphar.2016.06.046

Destreel G, Seutin V, Engel D. 2019. Subsaturation of the N-methyl-D-aspartate receptor glycine site allows the regulation of bursting activity in juvenile rat nigral dopamine neurons. Eur J Neurosci 50:3454–3471. doi:10.1111/ejn.14491

Drion G, Massotte L, Sepulchre R, Seutin V. 2011. How modeling can reconcile apparently discrepant experimental results: The case of pacemaking in dopaminergic neurons. PLoS Comput Biol 7:e1002050. doi:10.1371/journal.pcbi.1002050

Dufour MA, Woodhouse A, Amendola J, Goaillard JM. 2014. Non-linear developmental trajectory of electrical phenotype in rat substantia nigra pars compacta dopaminergic neurons. Elife 3. doi:10.7554/eLife.04059

Evans RC, Zhu M, Khaliq ZM. 2017. Dopamine inhibition differentially controls excitability of substantia nigra dopamine neuron subpopulations through T-type calcium channels. J Neurosci 37:3704–3720. doi:10.1523/JNEUROSCI.0117-17.2017

Farassat N, Costa Kauê Machado, Stovanovic S, Albert S, Kovacheva L, Shin J, Egger R, Somayaji M, Duvarci S, Schneider G, Roeper J. 2019. In vivo functional diversity of midbrain dopamine neurons within identified axonal projections. Elife 8. doi:10.7554/elife.48408

Galtieri DJ, Estep CM, Wokosin DL, Traynelis S, Surmeier DJ. 2017. Pedunculopontine glutamatergic neurons control spike patterning in substantia nigra dopaminergic neurons. Elife 6. doi:10.7554/eLife.30352

Giros B, Jaber M, Jones SR, Wightman RM, Caron MG. 1996. Hyperlocomotion and indifference to cocaine and amphetamine in mice lacking the dopamine transporter. Nature 379:606–612. doi:10.1038/379606a0

Gonon F, Bloch B. 1998. Kinetics and geometry of the excitatory dopaminergic transmission in the rat striatum in vivo. Adv Pharmacol 42:140–4. doi:10.1016/s1054-3589(08)60715-2

Grace AA, Bunney BS. 1984. The control of firing pattern in nigral dopamine neurons: Burst firing. J Neurosci 4:2877–2890. doi:10.1523/jneurosci.04-11-02877.1984

Guzman JN, Sánchez-Padilla J, Chan CS, Surmeier DJ. 2009. Robust Pacemaking in Substantia Nigra Dopaminergic Neurons. J Neurosci 29:11011–11019. doi:10.1523/JNEUROSCI.2519-09.2009

Held K, Voets T, Vriens J. 2016. Signature and Pathophysiology of Non-canonical Pores in Voltage-Dependent Cation Channels. Rev Physiol Biochem Pharmacol 170:67–99. doi:10.1007/112_2015_5003

Hong L, Pathak MM, Kim IH, Ta D, Tombola F. 2013. Voltage-Sensing Domain of Voltage-Gated Proton Channel Hv1 Shares Mechanism of Block with Pore Domains. Neuron 77:274–287. doi:10.1016/j.neuron.2012.11.013

Huang H, Tan BZ, Shen Y, Tao J, Jiang F, Sung YY, Ng CK, Raida M, Köhr G, Higuchi M, Fatemi-Shariatpanahi H, Harden B, Yue DT, Soong TW. 2012. RNA Editing of the IQ Domain in Cav1.3 Channels Modulates Their Ca2+-Dependent Inactivation. Neuron 73:304–316. doi:10.1016/j.neuron.2011.11.022

Ingram SL, Prasad BM, Amara SG. 2002. Dopamine transporter-mediated conductances increase excitability of midbrain dopamine neurons. Nat Neurosci 5:971–978. doi:10.1038/nn920

Jiang D, Gamal el-Din tamer M, Ing C, Lu P, Pomès R, Zheng N, Catterall WA. 2018. Structural basis for gating pore current in periodic paralysis. Nature 557:590–594. doi:10.1038/s41586-018-0120-4

Johnson SW, Seutin V, North RA. 1992. Burst firing in dopamine neurons induced by N-methyl-D-aspartate: Role of electrogenic sodium pump. Science 258:665–667. doi:10.1126/science.1329209

Kang Y, Kitai ST. 1993. Calcium spike underlying rhythmic firing in dopaminergic neurons of the rat substantia nigra. Neurosci Res 18:195–207. doi:10.1016/0168-0102(93)90055-U

Khalili-Araghi F, Tajkhorshid E, Roux B, Schulten K. 2012. Molecular dynamics investigation of the ω-current in the Kv1.2 voltage sensor domains. Biophys J 102:258–267. doi:10.1016/j.bpj.2011.10.057

Khaliq ZM, Bean BP. 2010. Pacemaking in Dopaminergic Ventral Tegmental Area Neurons: Depolarizing Drive from Background and Voltage-Dependent Sodium Conductances. J Neurosci 30:7401–7413. doi:10.1523/jneurosci.0143-10.2010

Khaliq ZM, Bean BP. 2008. Dynamic, Nonlinear Feedback Regulation of Slow Pacemaking by A-Type Potassium Current in Ventral Tegmental Area Neurons. J Neurosci 28:10905–10917. doi:10.1523/jneurosci.2237-08.2008

Lammel S, Hetzel A, Häckel O, Jones I, Liss B, Roeper J. 2008. Unique Properties of Mesoprefrontal Neurons within a Dual Mesocorticolimbic Dopamine System. Neuron 57:760–773. doi:10.1016/j.neuron.2008.01.022

Liss B, Roeper J. 2008. Individual dopamine midbrain neurons: Functional diversity and flexibility in health and disease. Brain Res Rev 58:314–321. doi:10.1016/j.brainresrev.2007.10.004

Margolis EB, Mitchell JM, Ishikawa J, Hjelmstad GO, Fields HL. 2008. Midbrain dopamine neurons: Projection target determines action potential duration and dopamine D2 receptor inhibition. J Neurosci 28:8908–8913. doi:10.1523/JNEUROSCI.1526-08.2008

Mason ER, Wu F, Patel RR, Xiao Y, Cannon SC, Cummins TR. 2019. Resurgent and gating pore currents induced by De Novo SCN2A epilepsy mutations. eNeuro 6:1–17. doi:10.1523/ENEURO.0141-19.2019

Mercuri NB, Bond A, Calabresi P, Stratta F, Stefani A, Bernardi G. 1994. Effects of dihydropyridine calcium antagonists on rat midbrain dopaminergic neurones. Br J Pharmacol 113:831–838. doi:10.1111/j.1476-5381.1994.tb17068.x

Moreau A, Gosselin-Badaroudine P, Chahine M. 2014. Molecular biology and biophysical properties of ion channel gating pores. Q Rev Biophys 47:364–388. doi:10.1017/S0033583514000109

Nedergaard S, Flatman JA, Engberg I. 1993. Nifedipine- and omega-conotoxin-sensitive Ca2+ conductances in guinea-pig substantia nigra pars compacta neurones. J Physiol 466:727–747. doi:10.1113/jphysiol.1993.sp019742

Neuhoff H, Neu A, Liss B, Roeper J. 2002. I(h) channels contribute to the different functional properties of identified dopaminergic subpopulations in the midbrain. J Neurosci 22:1290–302. doi:10.1523/JNEUROSCI.22-04-01290.2002

Nieoullon A. 2002. Dopamine and the regulation of cognition and attention. Prog Neurobiol 67:53–83. doi:10.1016/S0301-0082(02)00011-4

O’Leary T, Williams AH, Franci A, Marder E. 2014. Cell Types, Network Homeostasis, and Pathological Compensation from a Biologically Plausible Ion Channel Expression Model. Neuron 82:809–821. doi:10.1016/j.neuron.2014.04.002

Philippart F, Destreel G, Merino-Sepúlveda P, Henny P, Engel D, Seutin V. 2016. Differential Somatic Ca2+ Channel Profile in Midbrain Dopaminergic Neurons. J Neurosci 36:7234–7245. doi:10.1523/JNEUROSCI.0459-16.2016

Philippart F, Khaliq ZM. 2018. Gi/o protein-coupled receptors in dopamine neurons inhibit the sodium leak channel NALCN. Elife 7. doi:10.7554/eLife.40984

Poulin J-F, Zou J, Drouin-Ouellet J, Kim K-YA, Cicchetti F, Awatramani RB. 2014. Defining Midbrain Dopaminergic Neuron Diversity by Single-Cell Gene Expression Profiling. Cell Rep 9:930–943. doi:10.1016/j.celrep.2014.10.008

Puopolo M, Raviola E, Bean BP. 2007. Roles of Subthreshold Calcium Current and Sodium Current in Spontaneous Firing of Mouse Midbrain Dopamine Neurons. J Neurosci 27:645–656. doi:10.1523/JNEUROSCI.4341-06.2007

Richards CD, Shiroyama T, Kitai ST. 1997. Electrophysiological and immunocytochemical characterization of GABA and dopamine neurons in the substantia nigra of the rat. Neuroscience 80:545–557. doi:10.1016/S0306-4522(97)00093-6

Schultz W. 2007. Multiple Dopamine Functions at Different Time Courses. Annu Rev Neurosci 30:259–288. doi:10.1146/annurev.neuro.28.061604.135722

Seutin V, Engel D. 2010. Differences in Na+ conductance density and Na+ channel functional properties between dopamine and GABA neurons of the rat substantia nigra. J Neurophysiol 103:3099–114. doi:10.1152/jn.00513.2009

Seutin V, Massotte L, Renette MF, Dresse A. 2001. Evidence for a modulatory role of Ih on the firing of a subgroup of midbrain dopamine neurons. Neuroreport 12:255–258. doi:10.1097/00001756-200102120-00015

Sokolov S, Scheuer T, Catterall WA. 2010. Ion permeation and block of the gating pore in the voltage sensor of NaV1.4 channels with hypokalemic periodic paralysis mutations. J Gen Physiol 136:225–236. doi:10.1085/jgp.201010414

Sokolov S, Scheuer T, Catterall WA. 2007. Gating pore current in an inherited ion channelopathy. Nature 446:76–78. doi:10.1038/nature05598

Sokolov S, Scheuer T, Catterall WA. 2005. Ion Permeation through a Voltage-Sensitive Gating Pore in Brain Sodium Channels Having Voltage Sensor Mutations. Neuron 47:183–189. doi:10.1016/J.NEURON.2005.06.012

Sonders MS, Zhu SJ, Zahniser NR, Kavanaugh MP, Amara SG. 1997. Multiple ionic conductances of the human dopamine transporter: the actions of dopamine and psychostimulants. J Neurosci 17:960–74. doi:10.1523/JNEUROSCI.17-03-00960.1997

Starace DM, Bezanilla F. 2004. A proton pore in a potassium channel voltage sensor reveals a focused electric field. Nature 427:548–553. doi:10.1038/nature02270

Sun Y, Zhang XH, Selvaraj S, Sukumaran P, Lei S, Birnbaumer L, Brij X, Singh B. 2017. Inhibition of L-Type Ca 2 Channels by TRPC1-STIM1 Complex Is Essential for the Protection of Dopaminergic Neurons. J Neurosci 37:3364–3377. doi:10.1523/JNEUROSCI.3010-16.2017

Tarfa RA, Evans RC, Khaliq ZM. 2017. Enhanced sensitivity to hyperpolarizing inhibition in mesoaccumbal relative to nigrostriatal dopamine neuron subpopulations. J Neurosci 37:3311–3330. doi:10.1523/JNEUROSCI.2969-16.2017

Tepper JM, Damlama M, Trent F. 1994. Postnatal changes in the distribution and morphology of rat substantia nigra dopaminergic neurons. Neuroscience 60:469–477. doi:10.1016/0306-4522(94)90258-5

Tombola F, Pathak MM, Isacoff EY. 2005. Voltage-Sensing Arginines in a Potassium Channel Permeate and Occlude Cation-Selective Pores. Neuron 45:379–388. doi:10.1016/j.neuron.2004.12.047

Wise RA. 2004. Dopamine, learning and motivation. Nat Rev Neurosci 5:483–494. doi:10.1038/nrn1406

